# TTLL4 glutamyltransferase is a therapeutic target for NPM1-mutated acute myeloid leukemia

**DOI:** 10.1101/2025.04.07.647605

**Authors:** Alexandra Schurer, Humaira Ilyas, Maxim I. Maron, Subray Hegde, Melissa R. Leyden, Irene Roy, Rotila Hyka, Lucas Dada, Jeffrey Shabanowitz, Donald F. Hunt, Ellen Angeles, Victor Morell, Benjamin M. Lorton, Shira Glushakow-Smith, Daniel K. Borger, Yuxuan Wang, Linde Miles, Roger Belizaire, Seiya Kitamura, Kira Gritsman, David Shechter

**Author notes:** Contributed equally (Alexandra Schurer, Humaira Ilyas, Kira Gritsman, David Shechter).

## Abstract

NPM1-mutated acute myeloid leukemia (AML) is defined by aberrant cytoplasmic localization of the mutant NPM1c protein, and therapeutic strategies targeting this specific disease remain limited. Here, we identify TTLL4, a mono-glutamate glutamyltransferase, as a selective vulnerability in NPM1c AML. TTLL4 catalyzes post-translational hyper-glutamylation of NPM1c at E126, stabilizes its cytoplasmic localization and promotes a differentiation block in leukemic cells. Multiple genetic *TTLL4* inactivation approaches in human NPM1c-mutant cell lines reduce NPM1c glutamylation, trigger myeloid differentiation, and impair proliferation. Transcriptomic analyses show that *TTLL4* knockdown phenocopies NPM1c degradation and aligns with KMT2A and XPO1-targeted gene expression programs. Furthermore, *Ttll4* knockout significantly prolonged survival in an NPM1c/NRAS-driven mouse AML model and promoted differentiation. We identify a small molecule, EN7, that selectively inhibits TTLL4 and recapitulates these phenotypes in NPM1c^+^ cells. These findings identify glutamylation as a new axis of leukemic regulation and highlight TTLL4 as a druggable epigenetic regulator in NPM1c AML.

## Introduction

Acute myeloid leukemia (AML) is a devastating disease with a high relapse rate and a 5-year survival rate of just over 30 percent^1^. Nucleophosmin (*NPM1*) is one of the most frequently mutated genes in AML, with the most common mutation, called NPM1c, found in 30% of newly-diagnosed patients^2,3^. The rate of relapse and mortality of NPM1c AML is still high^4,5^. Therefore, there is an urgent need for therapeutic strategies to specifically target NPM1c.

NPM1 is a multi-functional nucleolar protein characterized by three major structural domains: the core domain, the intrinsically-disordered region (IDR), and the C-terminal three-helical bundle (3HB)^3^. The nucleolar localization signal (NoLS) within the 3HB is critical for NPM1 localization and function^2,3,6^. NPM1 mutations associated with AML result in the loss of the NoLS and the generation of a nuclear export signal (NES), resulting in mis-localization of mutant NPM1 to the cytoplasm; thus, the mutant protein is denoted as NPM1c^7^.

NPM1c AML is further characterized by increased expression of *HOXA/B* transcription factors and of their DNA-binding co-factor *MEIS1*, which is thought to be mediated by direct binding of NPM1c to chromatin through the second acidic stretch (A2) of the NPM1c IDR^8^. Aberrant *HOXA/B* and *MEIS1* expression in committed myeloid progenitors suppresses their differentiation and promotes self-renewal, transforming them into leukemic stem cells (LSCs)^9-11^. Although some progress has been made in understanding how NPM1c affects chromatin, there are still no approved therapeutic agents available to target these effects, and nothing is known about the contribution of post-translational modification of NPM1c to leukemogenesis^8,12^.

The structural domains of NPM1, particularly the acidic stretches of the IDR, are susceptible to various post translational modifications which play an important role in regulating NPM1 function. Tubulin tyrosine ligase-like 4 (TTLL4) is highly expressed in the bone marrow and catalyzes the post-translational glutamate-glutamylation of several histone chaperones, including NPM1 and its embryonic paralog, nucleoplasmin (NPM2)^13-15^. Notably, as *Ttll4* knockout mice have normal blood counts, TTLL4 does not appear to be required for normal hematopoiesis^14^. However, TTLL4 has been shown to play a role in cancer: TTLL4 overexpression causes abnormal proliferation of pancreatic ductal carcinoma cells through regulation of histone modifications and chromatin remodeling^16^.

Here, we find that NPM1c E126 is more heavily glutamylated by the TTLL4 glutamyltransferase than is wildtype NPM1 on E126 in the second acidic stretch. We demonstrate that inhibiting TTLL4 glutamyltransferase activity via shRNA knockdown or CRISPR-Cas9 deletion of *TTLL4* reduces proliferation of NPM1c AML cells and promotes myeloid differentiation with concomitant loss of self-renewal gene signatures, similar to changes that were observed with NPM1c degradation. Furthermore, loss of TTLL4 significantly prolongs survival in an NPM1c;NRAS-mutant mouse model of AML. Finally, we demonstrate that TTLL4 can be pharmacologically inhibited by a small molecule, causing loss of NPM1c glutamylation, reduced viability, and increased differentiation in NPM1c-mutant cells. These findings identify TTLL4-mediated NPM1c glutamylation as a novel mechanism of leukemic transformation and maintenance and reveal TTLL4 as a potential therapeutic target for NPM1c AML.

## Results

### NPM1c is preferentially glutamylated by TTLL4

The acidic stretches (A2 and A3) of the NPM1 and NPM1c C-terminal intrinsically-disordered regions (IDR) contain many glutamic and aspartic acid residues (**Figure 1A**). Glutamates in these acidic stretches are the substrates for glutamyltransferases, which post-translationally catalyze the reversible isopeptide addition of glutamic acids in chains of various lengths (**Figure 1B**)^13^. We and others previously showed that the mono-glutamyltransferase TTLL4 post-translationally glutamylates the histone chaperones Nap1, NPM1, and Npm2^13,15,17,18^. As the NPM1c mutation alters the conformation of the protein’s C-terminal domain and may therefore provide enhanced accessibility to the acidic stretches, we first sought to determine if mutant NPM1c is differentially glutamylated relative to wildtype protein. Immunoblot analysis of NPM1-wildtype OCI-AML2 and NPM1c-mutated OCI-AML3 and IMS-M2 cell lines revealed increased monoglutamylation of a 37 kDa band, specifically in mutant cells, which we confirmed by immunoprecipitation represented NPM1/NPM1c mono-glutamylation (**Figure 1C,D, Supplementary Figure S1A**). Notably, TTLL4 protein is expressed at similar levels among these cell lines, suggesting that hyper-glutamylation is specifically related to the presence of the NPM1c mutation (**Figure 1C**). Mass spectrometry analyses of cellular NPM1 and NPM1c protein confirmed hyper-glutamylation in the NPM1c-mutant cells (**Figure 1E, Supplementary Figure S1B**,**C**). This analysis also identified glutamate 126 (E126)—which resides in the A2 acidic stretch that is critical for NPM1c chromatin binding and orthologous to the histone binding domain we previously identified in Npm2—as the single mono-glutamylated residue of NPM1 and NPM1c in human AML cell lines (**Figure 1E**)^8^. Consistent with prior studies, our mass spectrometry analysis revealed abundant phosphorylation of the adjacent S125 in all three AML cell lines; interestingly, we found that for each E126 glutamylation, the adjacent S125 residue was also phos-phorylated (**Supplementary Figure S1C**)^18,19^.

**Figure 1.**
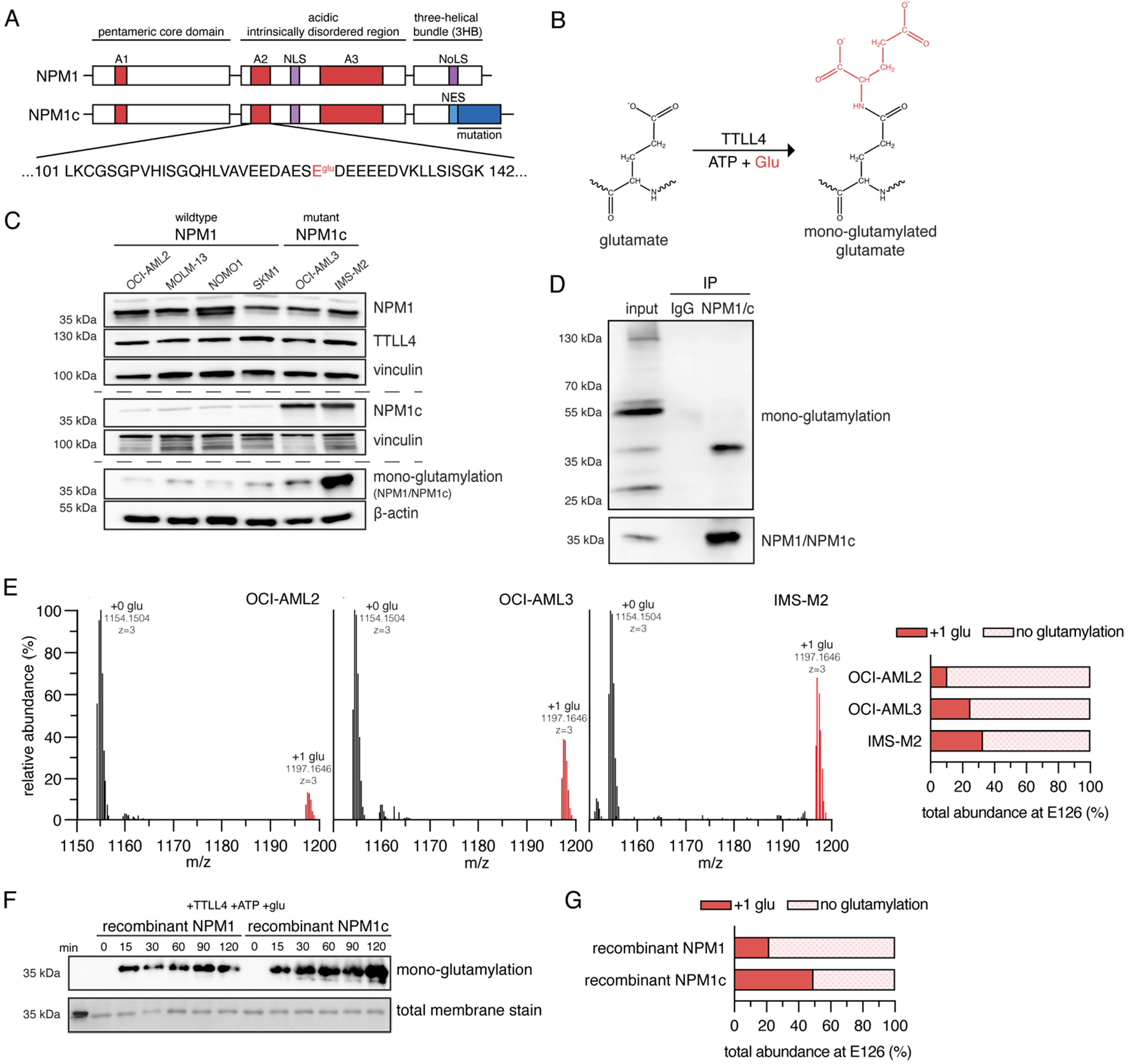
NPM1c is preferentially glutamylated by TTLL4. **(A)** Domain structures of wildtype NPM1 and mutated NPM1c (top). Sequence of NPM1/NPM1c peptides 101-142 in the second acidic stretch (A2) highlighting E126 as the main site of glutamylation (bottom). *A1, A2, A3 = acidic stretches; NES = nuclear export signal; NLS = nuclear localization signal; NoLS = nucleolar localization signal*. **(B)** Schematic of the glutamylation reaction. The TTLL4 glutamyltransferase enzyme post-translationally adds a glutamic acid residue to a protein glutamate. **(C)** Immunoblot analysis of NPM1/NPM1c, TTLL4, and mono-glutamylation levels in NPM1-wildtype and NPM1c-mutant AML cell lines. Vinculin and β-actin were used as loading controls. **(D)** Immunoblot stained with mono-glutamylation and total NPM1/NPM1c antibodies depicting bands at ∼37 kDa corresponding to glutamylated NPM1/NPM1c proteins isolated from OCI-AML3 using immunoprecipitation. This cellular protein was used for mass spectrometric analysis. **(E)** Mass spectrometry (MS1) of glutamylation of amino-affinity purified NPM1 or NPM1c E126 in the NPM1-wildtype OCI-AML2 cell line and the NPM1c-mutated OCI-AML3 and IMS-M2 cell lines. Peaks are labeled with the C12 monoisotopic mass of each peptide (left). Quantification of relative abundance of E126 mono-glutamylation across the three cell lines (right). **(F)** Immunoblot of recombinant NPM1 and NPM1c incubated with TTLL4 enzyme, ATP, and glutamate for 0, 15, 30, 60, 90, and 120 minutes probed with mono-glutamylation antibody and Direct Blue 71 to measure total transferred protein content. (G) Quantification of the relative abundance by mass spectrometry (MS1) of E126 mono-glutamylation of recombinant NPM1 and NPM1c.

To determine if NPM1c is a better substrate for TTLL4 than wildtype NPM1, we treated purified recombinant NPM1 and NPM1c with TTLL4 enzyme catalytic domain *in vitro* (**Supplementary Figure S1D**,**E**). Consistent with our findings in AML cell lines, we observed by immunoblot analysis that recombinant NPM1c, compared to wildtype NPM1, is a better substrate for glutamylation (**Figure 1F**). Mass spectrometry analysis also confirmed that NPM1c is more abundantly mono-glutamylated at E126 than is the wildtype protein (**Figure 1G, Supplementary Figure S1C**,**F**). Overall, these findings suggest that NPM1c is a preferred substrate of TTLL4.

### Loss of TTLL4 impairs proliferation and promotes differentiation in NPM1-mutated AML cells

To determine if TTLL4-mediated NPM1c glutamylation contributes to maintenance of the AML cell pheno-type, particularly increased proliferation and self-renewal, we used multiple approaches to genetically inactivate *TTLL4* with resulting loss of NPM1 and NPM1c glutamylation in the NPM1c-mutated OCI-AML3 human cell line^20^. First, we found that CRISPR-Cas9 deletion of *TTLL4* using two different sgRNAs (sgTTLL4) led to a specific reduction of mono-glutamylation on NPM1/NPM1c (**Figure 2A**). Expression of two different *TTLL4*-targeting sgRNAs (sgTTLL4) inhibited OCI-AML3 cell proliferation, with the greatest decrease in proliferation compared to non-targeting (sgNT) controls observed at 6 days after TTLL4 loss (**Figure 2B**). To examine the clonogenic potential of *TTLL4* knockout in OCI-AML3 cells, we performed colony-forming unit (CFU) assays in methylcellulose media supplemented with myeloid growth factors. We observed a significant decrease in the number of CFU formed by sgTTLL4 cells compared to sgNT controls (**Figure 2C**). These findings support our hypothesis that TTLL4-mediated glutamylation contributes to the proliferation and clonogenic activity of NPM1c-mutated AML cells.

**Figure 2.**
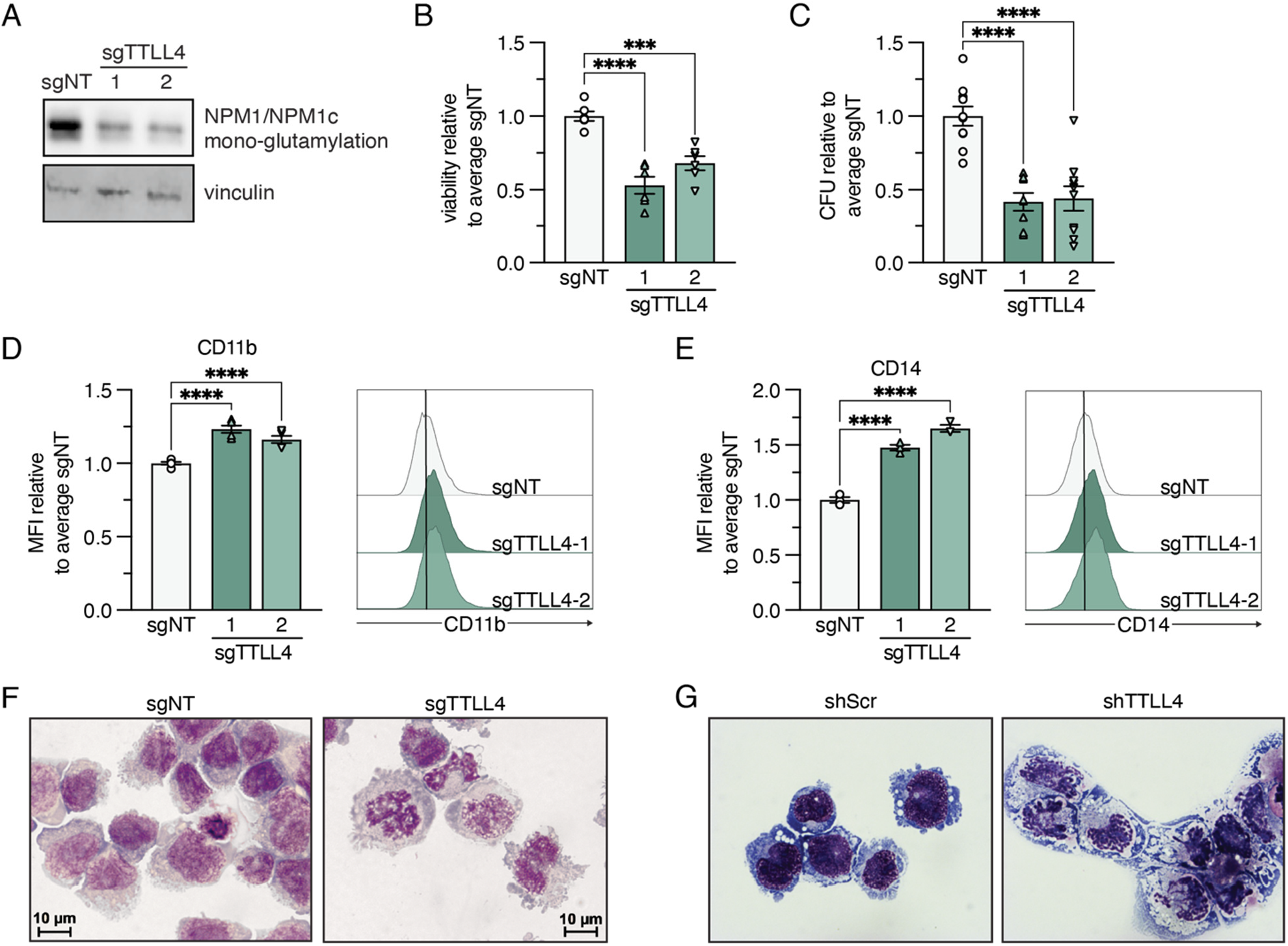
Loss of TTLL4 impairs proliferation and promotes myeloid differentiation of NPM1c-mutated AML cell lines. (A-G) Assays of the NPM1c-mutant OCI-AML3 human AML cell line with or without CRISPR-Cas9 deletion of *TTLL4* by expression of *TTLL4* (sgTTLL4) or non-targeting (sgNT) control sgRNAs. **(A)** Validation of CRISPR-Cas9 *TTLL4* deletion in sgTTLL4-transduced cells by immunoblot analysis of mono-glutamylation of NPM1 and NPM1c (∼37 kDa) with vinculin loading control. (B-C) NPM1c-mutant cell proliferation measured by **(B)** normalized cell viability at day 6 using CellTiter-Glo^®^ proliferation assays and **(C)** normalized number of colony-forming units (CFU) formed from 5 x10^3^ cells by day 7 in OCI-AML3 cells. **(D-E)** Normalized median fluorescent intensity (MFI) of **(D)** CD11b and **(E)** CD14 expression at day 8 after CRISPR-Cas9. Representative histograms for one experimental replicate are shown. **(F)** Wright-Giemsa staining of cytospins of OCI-AML3 cells at day 8. Representative images from one experimental replicate are shown. **(G)** Wright-Giemsa staining of cytospins of NPM1c-FKBP12^F36V^ (NPM1c-degron) OCI-AML3 cells 8 days after doxycycline-induction of *TTLL4* (shTTLL4) or scrambled (shScr) shRNAs. Representative images from one experimental replicate are shown. Error bars indicate mean ± SEM; ***p ≤0.001, ****p ≤0.0001; ordinary one-way ANOVA.

We then investigated whether TTLL4 loss induces differentiation in NPM1c cells. Using flow cytometry with the myeloid markers CD11b and CD14, our analysis demonstrated that *TTLL4* deletion significantly promotes myeloid differentiation by day 8, as CRISPR-Cas9-mediated deletion using two different *TTLL4*-targeting sgRNAs significantly increased CD11b and CD14 expression in OCI-AML3 cells (**Figure 2D,E**). Myeloid differentiation was also evident by analysis of cellular morphology at this time point, which showed a decreased nuclear to cytoplasmic ratio and myeloid nuclear morphology after loss of TTLL4 in OCI-AML3 cells (**Figure 2F**).

Consistent with our findings in the *TTLL4*-deleted cells, we found that knockdown of *TTLL4* expression using two different shRNAs (shTTLL4) also significantly reduced NPM1c glutamylation and inhibits OCI-AML3 cell proliferation relative to scrambled (shScr) controls (**Supplementary Figure S2A**,**B)**. shTTLL4 cells also developed morphological changes consistent with myeloid differentiation by 8 days after *TTLL4* knockdown (**Figure 2G**). Altogether, these findings support our hypothesis that TTLL4 is a regulator of leukemic transformation in hematopoietic cells.

### TTLL4 loss re-localizes NPM1/NPM1c to the nucleus and promotes myeloid gene expression

To determine if TTLL4 loss phenocopies NPM1c loss at the gene expression level, we performed RNA sequencing in NPM1c-degron OCI-AML3 cells with either doxycycline-inducible *TTLL4* shRNA knockdown (TTLL4*kd*) or dTAG-13-inducible degradation of the NPM1c-FKBP12^F36V^ fusion protein (NPM1c-dTag)^21^. This approach allows for more precise temporal control of glutamylation loss and endogenous NPM1c degradation (**Supplementary Figure S2A, Supplementary Figure S3A**)^21^. We observed that *TTLL4* knockdown upregulated the expression of multiple myeloid genes such as *S100A9* and *CD226* without downregulating expression of *MEIS1* or *HOX* genes (**Figure 3A**). However, several other oncogenes were found to be downregulated with loss of TTLL4, including *GATA2* and *WT1* (**Figure 3A**). Degradation of NPM1c without the loss of TTLL4 (NPM1c-dTag) produced a comparable pattern of gene expression changes, as seen in the correlation plot, supporting that TTLL4-mediated glutamylation promotes leukemogenesis in part through an NPM1c-dependent mechanism (**Figure 3B, Supplementary Figure S3B**). We did observe that many downregulated and upregulated genes did not overlap between the TTLL4*kd* and NPM1c-dTag conditions (**Supplementary Figure S3C**,**D**). The inverse regulation of genes such as *IL18R* and *CDK2NA* between the two conditions (**Supplementary Figure S3B**) suggests that *TTLL4* knockdown may affect additional NPM1c-independent processes or that glutamylation is dispensable for certain NPM1c-driven transformation mechanisms.

**Figure 3.**
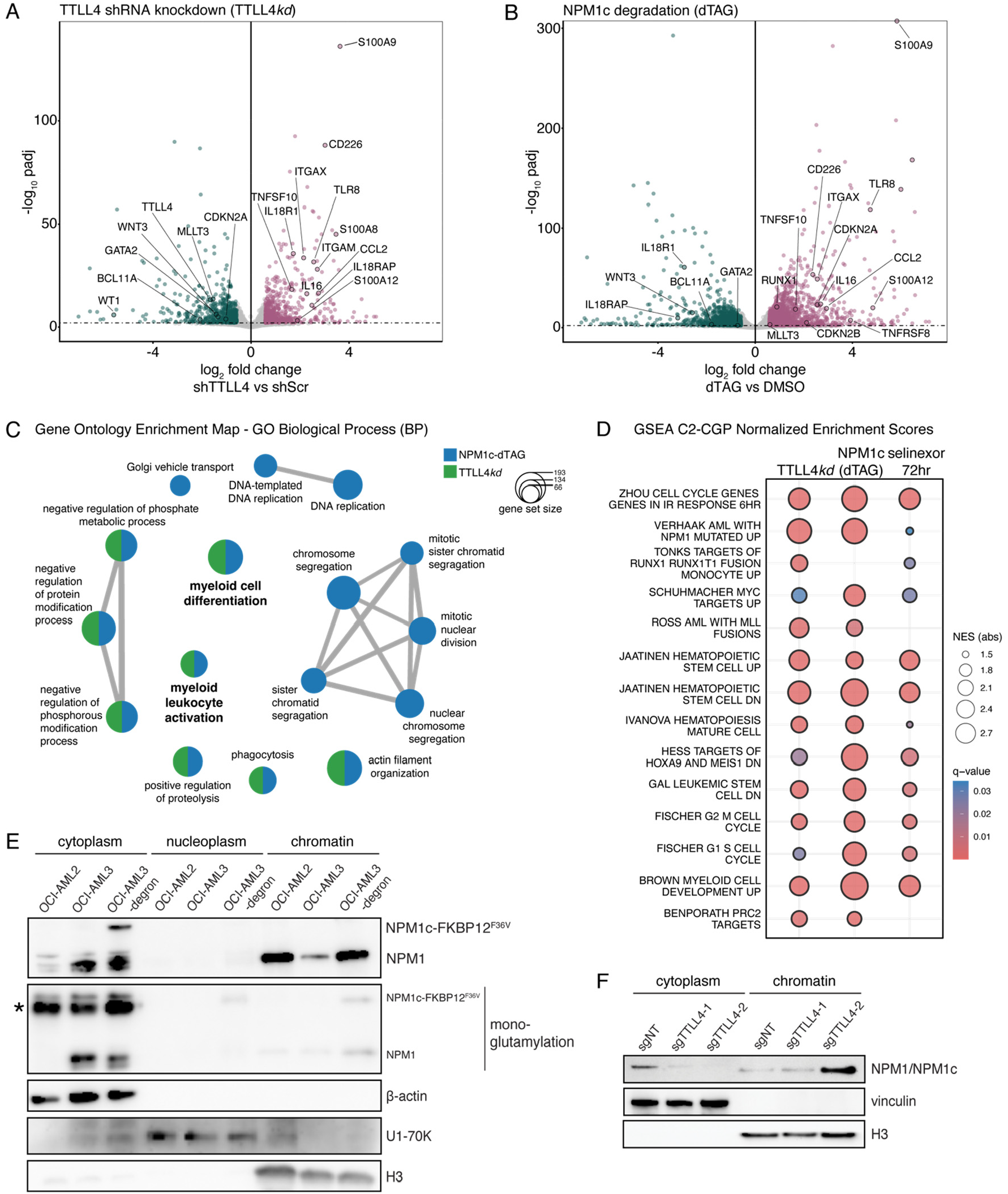
Loss of TTLL4 impairs proliferation and promotes myeloid differentiation of NPM1c-mutated AML cell lines. (A) Volcano plot of altered genes revealed by mRNA-seq of OCI-AML3 NPM1c-degron cells with TTLL4 shRNA-mediated knockdown (TTLL4kd) compared to scrambled (shScr) controls. Representative oncogenes and differentiation markers are indicated. (B) Volcano plot of dTAG-treated OCI-AML3 NPM1c-degron cell mRNA-seq compared to DMSO controls with relevant genes highlighted. (C) ClusterProfiler over-representation analysis enrichment map for NPM1c-dTAG (blue) and TTLL4kd (green) Gene Ontology (GO) Biological Process (BP) terms. (D) Gene Set Enrichment Analysis (GSEA) dot plot of relevant AML and cell cycle terms for TTLL4kd, NPM1c dTAG treatment, and publicly available selinexor-treated OCI-AML3 cell data. (E) Immunoblot analysis of fractionated sub-cellular compartments of NPM1-wildtype OCI-AML2 and NPM1c-mutated OCI-AML3 and OCI-AML3 NPM1c-degron cells. Fractions were probed with total NPM1/NPM1c antibody and mono-glutamylation antibody with β-actin, U1-70K, and H3 antibodies as loading controls for the cytoplasmic, nucleoplasmic, and chromatin fractions, respectively. * indicates an additional (non-NPM1c) glutamylated protein at ∼50 kDa that we have not identified but is likely β-tubulin, a known substrate of TTLL4. (F) Immunoblot analysis of fractionated cytoplasmic and chromatin compartments of OCI-AML3 cells with or without CRISPR-dCas9 knockdown of TTLL4 after expression of TTLL4 (sgTTLL4) or non-targeting (sgNT) control sgRNAs.

Gene ontology (GO) over-enrichment analysis shows that many of the same biological processes and molecular functions are enriched in both TTLL4*kd* and NPM1c-dTag cells, including those related to myeloid cell differentiation and myeloid leukocyte activation (**Figure 3C, Supplementary Figure S3E**). However, we found that only NPM1c-dTag cells had over-representation of gene signatures related to enriched for mitotic cell division and DNA replication, despite established roles for both glutamylation and NPM1 in mitosis (**Figure 3C, Supplementary Figure S3E**)^22-25^. These observations support the existence of glutamylation-independent mechanisms of NPM1c leukemogenesis.

Gene Set Enrichment Analysis (GSEA) for selected AML gene sets revealed that both *TTLL4* knockdown and NPM1c degradation resulted in positive enrichment of myeloid differentiation and cell cycle-related gene signatures relative to controls (**Figure 3D**). Interestingly, we also observed negative enrichment of self-renewal gene sets and leukemogenic gene sets in both groups, despite neither *HOXA9* nor *MEIS1* genes being downregulated in TTLL4*kd* cells (**Figure 3D**).

To test if transcriptomic changes in TTLL4*kd* and NPM1c-dTag cells significantly overlapped with those induced by inhibition of the KMT2A complex, we used GeneOverlap to perform a Fisher Exact Test between these datasets. We observed significant overlap (odds ratios) of both TTLL4*kd* and NPM1c-dTag cells with inhibition of various components of the KMT2A complex and its interactors, including menin, ENL, KAT6A, or the BET protein BRD4 (**Supplementary Figure S3F**)^26-28^. The menin-KMT2A complex has been identified as a key mediator of NPM1c chromatin binding and leukemogenic gene transcription; therefore, our findings suggest that TTLL4-mediated glutamylation has a plausible role in modulating the KMT2A-mediated NPM1c transcriptional response^8,26^.

The transcriptomic changes observed in both the TTLL4*kd* and NPM1c-dTag conditions were also similar to those identified in OCI-AML3 cells treated with the XPO1-inhibitor selinexor, which promotes myeloid differentiation by re-localizing NPM1c from the cytoplasm to the nucleus (**Figure 3D, Supplementary Figure S3F**)^21^. To test if TTLL4 influences NPM1/NPM1c localization, we performed immunoblotting after sub-cellular fractionation of OCI-AML3 cells. This revealed that glutamylation is mostly restricted to cytoplasmic NPM1 and mutant NPM1c, with very little glutamylated protein being observed in the chromatin fraction (**Figure 3E**). To determine if glutamylation contributes to the cytoplasmic mis-localization of NPM1c, we performed immunoblotting after sub-cellular fractionation of OCI-AML3 cells with CRISPR-dCas9 knockdown of *TTLL4*, which phenocopied CRISPR-Cas9 deletion of *TTLL4* (**Supplementary Figure S3G-I**). Notably, we observed depletion of total NPM1/NPM1c protein in the cytoplasmic fraction of sgTTLL4 cells and concomitant increases in the chromatin fraction. These observations support the hypothesis that NPM1/NPM1c re-localize to the nucleus upon loss of *TTLL4* expression (**Figure 3F**).

### TTLL4 is necessary for clonogenic activity of NPM1c mutant mouse cells

To further characterize the effects of TTLL4 loss in NPM1c hematopoietic cells, we performed *ex vivo* CRISPR-Cas9 deletion of *Ttll4* in c-Kit^+^ immature mouse bone marrow cells harvested from *Npm1*^*frt-cA/+*^*;R26*^*FlpoER*^ mice expressing the *Npm1*^*cA*^ mutant allele or wildtype C57BL/6 controls^29^. Heterozygous expression of the *Npm1*^*cA*^ mutant allele after tamoxifen induction of FLPo recombinase was validated by PCR, and *Ttll4* deletion after CRISPR was confirmed by immunoblotting (**Supplementary Figure S4A**,**B**)^29^. *Npm1*^*cA/+*^ mice were previously reported to develop a mild expansion of myeloid cells, but do not develop AML without additional mutations^29^. Consistent with our results in OCI-AML3 cells, we found that NPM1c-mutant mouse cells had impaired colony-forming ability upon loss of TTLL4 **(Figure 4A**). In contrast, there was no such effect in wildtype cells, suggesting that the effects of *Ttll4* deletion in hematopoietic cells could be selective for NPM1-mutated cells (**Figure 4B**).

**Figure 4.**
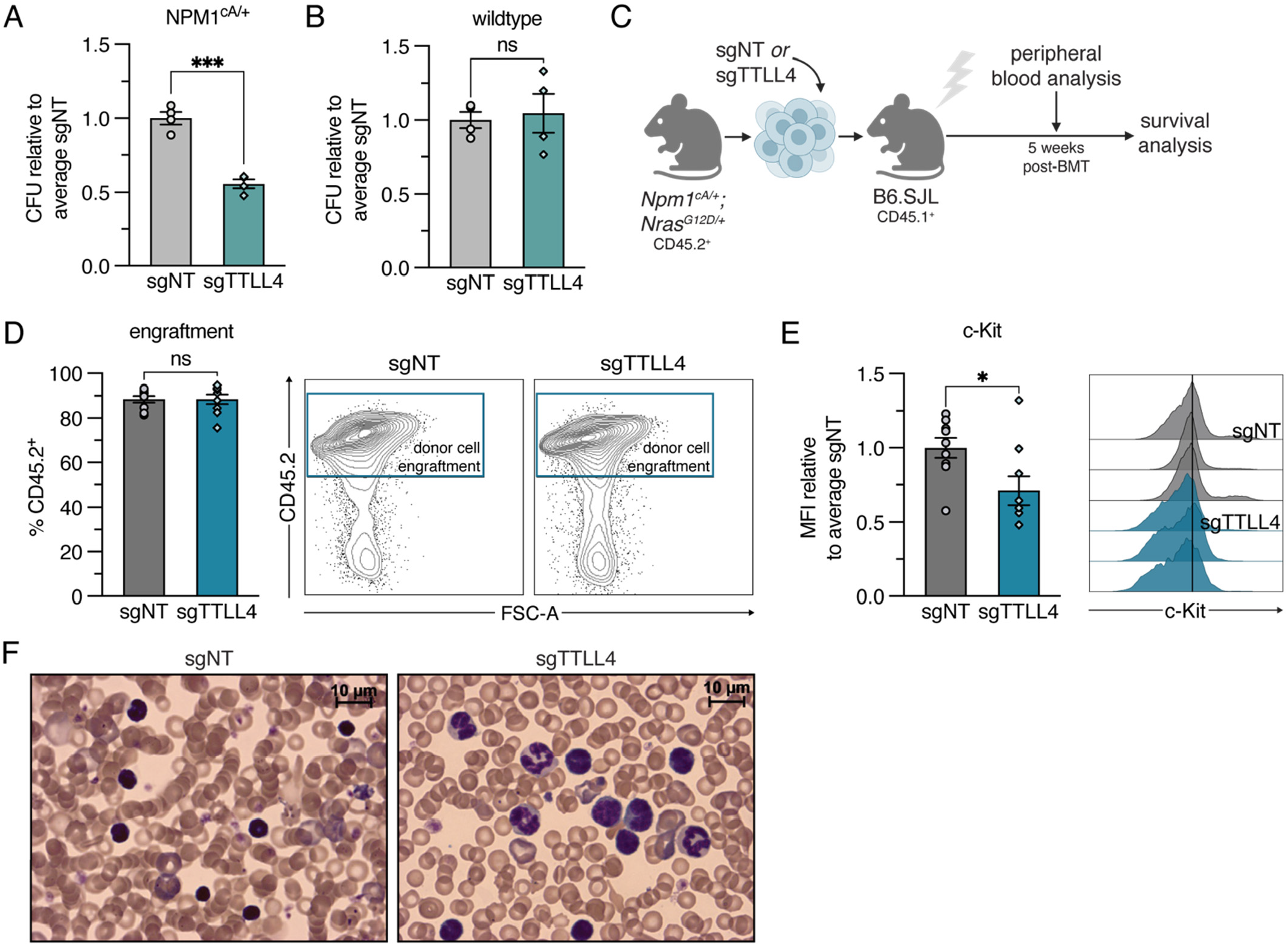
TTLL4 is necessary for leukemia maintenance in mouse models of NPM1c AML. **(A)** Normalized number of colony-forming units (CFU) formed by day 7 from 3 x10^3^ c-Kit^+^ cells harvested from *Npm1*^*cA/+*^ mouse bone marrow after *Ttll4* CRISPR editing. **(B)** Normalized number of colony-forming units (CFU) formed by day 7 from 3 x10^3^ c-Kit^+^ cells harvested from wildtype (C57/Bl6) mouse bone marrow after *Ttll4* CRISPR editing. Experiments in (A) and (B) were performed three times with similar results. **(C)** Experimental design for primary transplant assays of *Ttll4* CRISPR-Cas9 knockout in the *Npm1*^*cA/+*^;*Nras*^*G12D/+*^ model of AML. Whole bone marrow cells were harvested from leukemic CD45.2^+^ *Npm1*^*cA/+*^;*Nras*^*G12D/+*^ primary transplant-recipient mice, electroporated with sgNT or sgTTLL4 Cas9-ribonucleoproteins (RNPs), and transplanted into sub-lethally irradiated CD45.1^+^ B6.SJL recipient mice (n = 9). **(D)** Engraftment of donor cells was measured by flow cytometry for CD45.2^+^ cells in peripheral blood collected at 5 weeks post-transplantation. Representative flow plots for one mouse per group are shown. **(E)** Normalized median fluorescent intensity (MFI) of c-Kit expression in CD45.2^+^ peripheral blood cells at 5 weeks. Representative histograms from 3 mice per group are shown. **(F)** Wright-Giemsa staining of peripheral blood smears at 5 weeks. Representative images from one mouse per group are shown. Normalized data is presented as the ratio of individual replicates over the average of all sgNT replicates. Error bars indicate mean ± SEM; *p ≤0.05, ***p ≤0.001; unpaired t-test with Welch’s correction.

### Loss of TTLL4 prolongs survival and promotes myeloid differentiation in a mouse model of NPM1c AML

To better model the *Ttll4* deletion phenotype in an NPM1c AML model *in vivo*, we performed CRISPR-Cas9 deletion of *Ttll4* in leukemic bone marrow harboring both NPM1^cA^ and NRAS^G12D^ mutations^11^. CD45.2^+^ *Npm1*^*cA/+*^;*Nras*^*G12D/+*^ whole bone marrow was harvested from a leukemic donor mouse after disease onset and electroporated with sgNT or sgTTLL4 with Cas9-ribonu-cleoproteins (RNPs) (**Supplementary Figure S4C**). Electroporated cells were then transplanted into sub-lethally irradiated CD45.1^+^ recipient mice (**Figure 4C**). We confirmed engraftment of CD45.2^+^ leukemic cells by peripheral blood flow cytometry (**Figure 4D**). Analysis of peripheral blood collected at five weeks post-transplantation revealed that white blood cell counts (WBC) were elevated in both groups (**Supplementary Figure S4D**). However, most mice were not yet symptomatic, so this analysis represented a pre-clinical disease timepoint. We found that while there was no difference in the percent engraftment of leukemic *Npm1*^*cA/+*^;*Nras*^*G12D/+*^ cells in the peripheral blood, the fraction of leukemic cells expressing the immature marker c-Kit was decreased in the sgTTLL4 group compared to non-targeting controls, providing evidence of differentiation *in vivo* at a pre-clinical disease timepoint (**Figure 4D,E**). This is consistent with the presence of cells with evidence of myeloid maturation in the peripheral blood of sgTTLL4 mice (**Figure 4F**). These findings suggest that *Ttll4* deletion promotes myeloid maturation in NPM1c AML *in vivo*.

We found that recipients of sgTTLL4-electroporated *Npm1*^*cA/+*^;*Nras*^*G12D/+*^ leukemic cells had significantly prolonged survival compared with mice in the sgNT group before succumbing to AML (median survival of 59 days in sgTTLL4 group and 36 days in sgNT group, p = 0.008) (**Figure 5A**). We did not observe significant differences in the blood counts between the two groups at the time of death, and editing was confirmed by Sanger sequencing in all leukemic mice in the sgTTLL4 group (**Supplementary Figure S5A**). Consistent with the decrease in the c-Kit^+^ population observed in the peripheral blood at the pre-disease timepoint, we observed that bone marrow from the sgTTLL4 mice had a significantly decreased frequency of the LK cell (Lineage^-^ Sca1^-^ c-Kit^+^) population, which represents myeloid progenitors (**Figure 5B**). However, we did not observe any significant differences in the frequency of LSK (Lineage^-^ Sca1^+^ c-Kit^+^) immature cells (**Supplementary Figure S5B**). Notably, prior characterization of this mouse model revealed that *Npm1*^*cA/+*^;*Nras*^*G12D/+*^ mice have a significant expansion of the LK population^11^. Therefore, our data suggests that deletion of *Ttll4* at least in part reverses the dysregulation of hematopoiesis in these mice.

**Figure 5.**
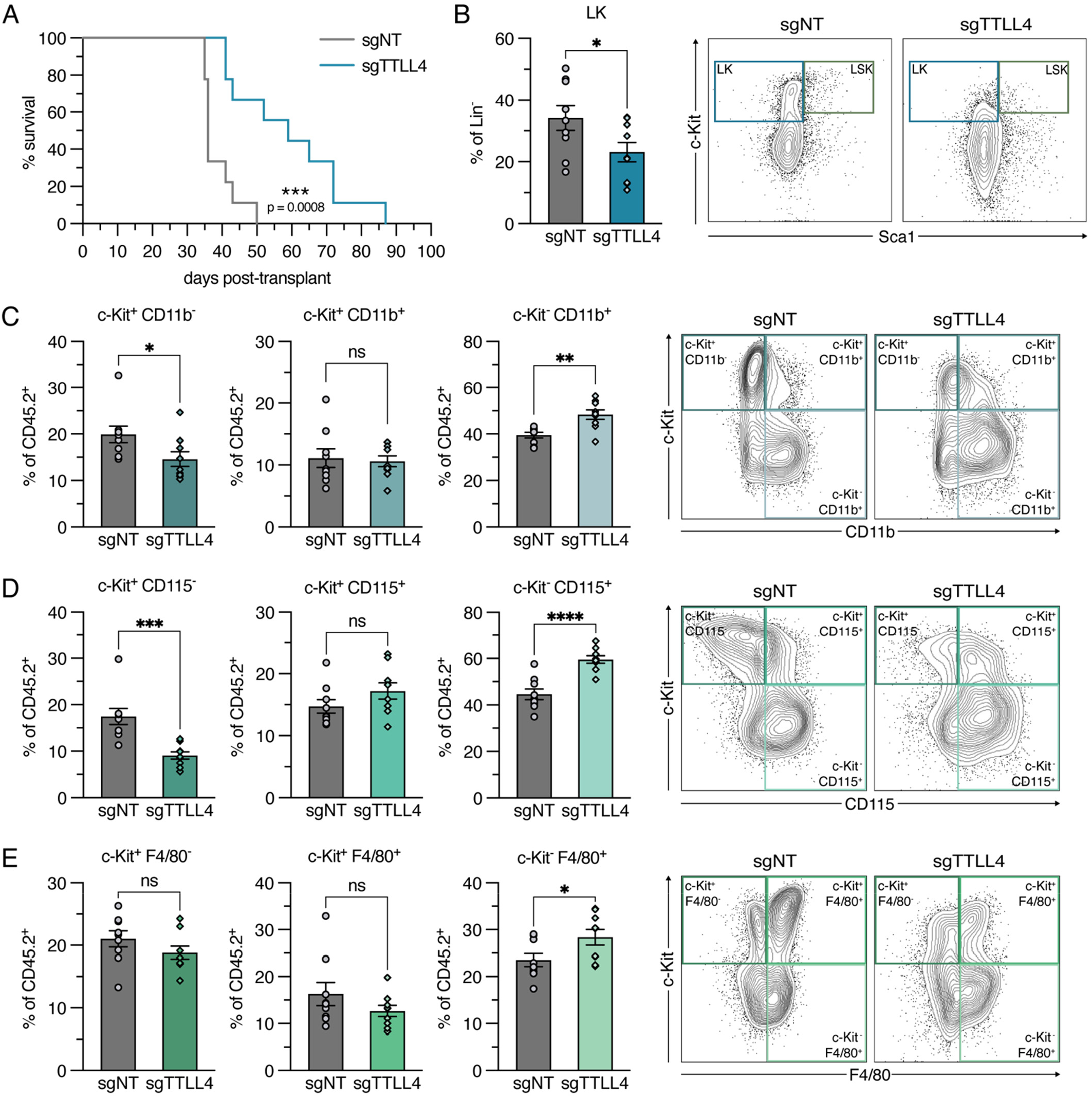
Loss of TTLL4 prolongs survival and promotes myeloid differentiation in NPM1c AML with a cooperating NRAS mutation. **(A)** Kaplan-Meier survival curves of mice transplanted with *Npm1*^*cA/+*^;*Nras*^*G12D/+*^ leukemic bone marrow cells electroporated with sgNT or sgTTLL4 RNPs (n = 9 per group). **(B)** Quantification of flow cytometry analysis of Lin^-^ Sca1^-^ c-Kit^+^ (LK) myeloid progenitors in control (sgNT) or *Ttll4*-deleted (sgTTLL4) CD45.2^+^ bone marrow collected at time of death. Representative flow plots of LK and LSK (Lin^-^ Sca1^+^ c-Kit^+^) populations for one mouse per group are shown. (C-E) Quantification of flow cytometry analysis of expression of the immature cell marker c-Kit against the myeloid differentiation markers **(C)** CD11b, **(D)** CD115, and **(E)** F4/80 in CD45.2^+^ bone marrow collected at time of death. Representative flow plots from one mouse per group are shown. Error bars indicate mean ± SEM; *p ≤0.05, **p ≤0.01, ***p ≤0.001, ****p ≤0.0001; Log-rank Mantel-Cox test (A), unpaired t-test with Welch’s correction (B-E).

Consistent with the diminished hematopoietic stem and progenitor cell populations, we observed a decrease in c-Kit-positive populations and an increase in the c-Kit-negative CD11b^+^, CD115^+^, and F4/80^+^ myeloid populations in the sgTTLL4 group, consistent with myeloid differentiation of leukemic blasts (**Figure 5C-E**). Together, this indicates an important role for TTLL4 in NPM1c-mediated leukemogenesis *in vivo*, likely through inhibition of NPM1c-mutant cell differentiation.

### TTLL4 can be pharmacologically targeted in NPM1c AML

To facilitate the identification of small-molecule inhibitors for TTLL4, we first constructed a structural model of human TTLL4 bound to a previously identified TTLL4 substrate, *Xenopus tropicalis* Nap1L4, using AlphaFold-Multimer (**Figure 6A**)^17,30^. This modeling revealed a putative acidic peptide substrate binding pocket and showed critical residues within the TTLL4 catalytic domain (M551-G1078) necessary for activity, including ATP-binding residues (K721, R727-728, K762-764, T809-810), glutamate-binding residues (R727, R788, K924), and magnesium-binding residues (D893, E906, N908) (**Supplementary Figure S6A**). Based on homology with TTLL6 (PDB IDs: 6VZW, 6VZT), we modeled into TTLL4 the ATP substrate and the TTLL6 “initiation” analog 2TI that was previously found to be a glutamate transition state mimic^31^. Consistent with our identification of the true TTLL4 catalytic site, a modeled NAP1L4 peptide and a modelled NPM1 A2 acidic stretch peptide substrate glutamate occupied the same space as 2TI. To test TTLL4 activity, we employed a dot-blot mono-glutamylation antibody-based assay using either NAP1L4 or NPM1c as a protein substrate. We also experimentally validated the TTLL4 structural model experimentally by testing an E905A TTLL4 mutant, which was deficient in glutamyl-transferase activity (**Supplementary Figure S6B**).

**Figure 6.**
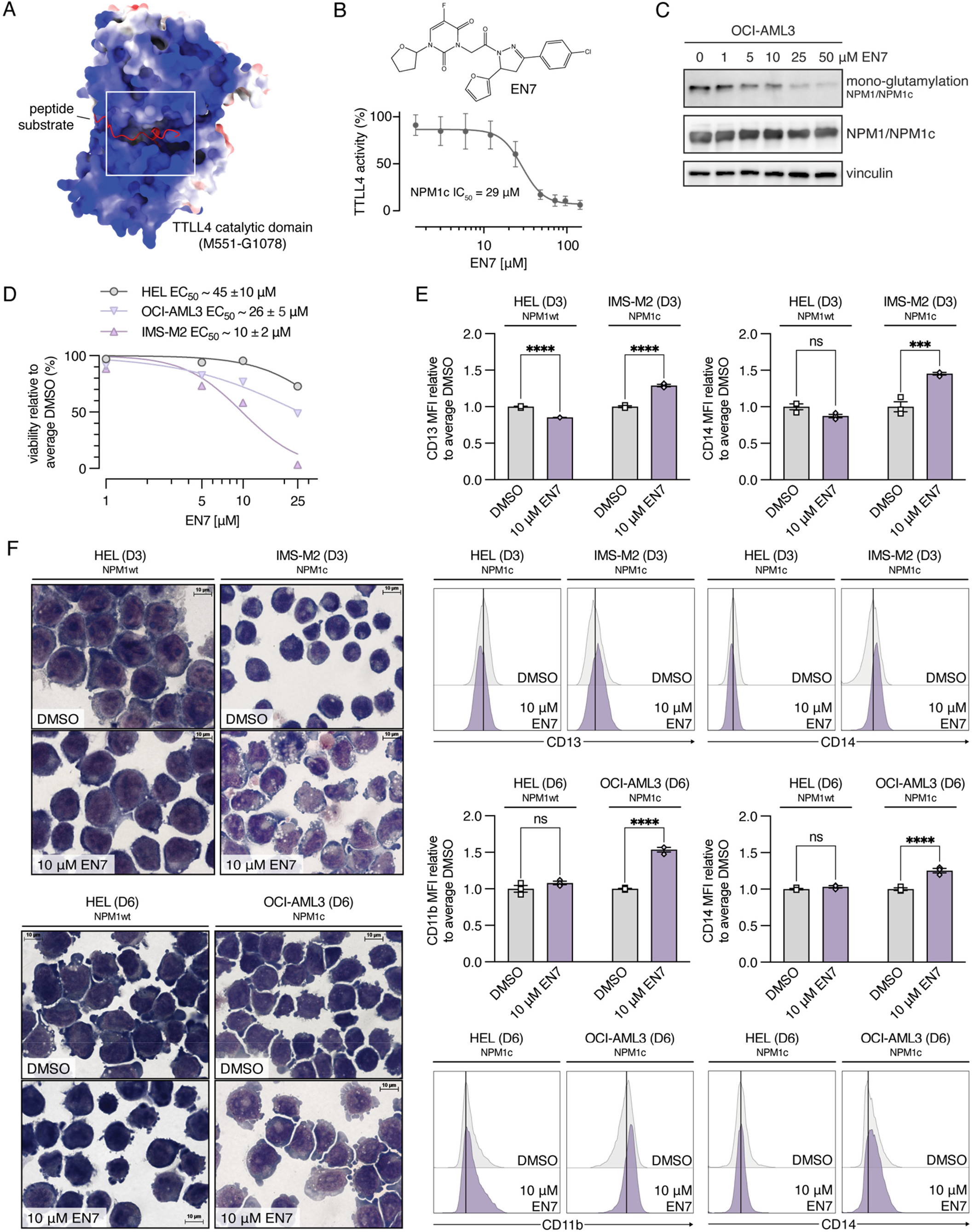
TTLL4 can be pharmacologically targeted in NPM1c AML. **(A)** Structural model of TTLL4 catalytic domain in complex with an acidic IDR substrate and ATP revealed a deep binding pocket. **(B)** EN7 inhibited recombinant TTLL4 glutamyltransferase activity towards NPM1c with an *in vitro* IC_50_ of 29 μM. **(C)** Immunoblot validation of inhibition of TTLL4-mediated mono-glutamylation in OCI-AML3 cells on day 3 of treatment with EN7 or DMSO control. Total NPM1/NPM1c expression and vinculin were used as loading controls. **(D)** Dose response curve on day 6 of treatment of NPM1-wildtype (HEL) and NPM1c-mutated (IMS-M2, OCI-AML3) human AML cell lines with EN7. EN7 inhibited HEL cell proliferation with an EC_50_ of ∼45 ± 10 μM and inhibited OCI-AML3 and IMS-M2 cell proliferation with EC_50_ values of ∼ 26 ± 5 μM and ∼10 ± 2 μM, respectively. **(E)** Normalized median fluorescent intensity (MFI) of CD13 and CD14 expression in HEL and IMS-M2 cells on day 3 of treatment with 10 μM EN7 or DMSO control (top). Normalized median fluorescent intensity (MFI) of CD11b and CD14 expression in HEL and OCI-AML3 cells on day 6 of treatment with 10 μM EN7 or DMSO control (bottom). Representative histograms for one replicate per cell line are shown. **(F)** Wright-Giemsa staining of cytospins of HEL and IMS-M2 cells on day 3 of treatment with 10 μM EN7 or DMSO control (top). Wright-Giemsa staining of cytospins of HEL and OCI-AML3 cells on day 6 of treatment with 10 μM EN7 or DMSO control (bottom). Representative images from one experimental replicate are shown. Error bars indicate mean ± SEM; ***p ≤0.001, ****p ≤0.0001; ordinary two-way ANOVA.

To identify TTLL4 small molecule inhibitors, we performed ligand-based (2TI and ATP) and structure-based virtual screening approaches; we also screened the Enamine nucleoside mimetic library. Two compounds, EN7 (from the nucleoside mimetic library screen) and TM42 (from the structure-based virtual screen), were identified as bona fide hits with reproducible TTLL4 inhibition, acceptable physicochemical properties, and reliable docking, although only EN7 remained soluble in tissue culture media (**Supplementary Figure S6C-D**). EN7 inhibited recombinant TTLL4 catalytic domain enzymatic activity toward NPM1c with an *in vitro* IC_50_ of 29 μM (**Figure 6B**), representing the first-in-class small molecule TTLL4 inhibitor.

Next, we sought to determine if pharmacologic inhibition of TTLL4 phenocopies the proliferation defects and differentiation phenotype associated with genetic deletion of TTLL4 in NPM1c AML. Loss of glutamylation of NPM1 and NPM1c was detectable with EN7 doses as low as 5 μM in OCI-AML3 cells after three days of treatment (**Figure 6C**). However, we found that treatment with doses as low as 1 μM specifically diminished cellular proliferation of two different NPM1c-mutated human AML cell lines, IMS-M2 and OCI-AML3, while having minimal effect on the proliferation of NPM1-wildtype HEL cells (**Figure 6D, Supplementary Figure S6E**). Specifically, EN7 decreased cell viability at day 6 with EC_50_ values of ∼ 26 ± 5 μM and ∼ 10 ± 2 μM in OCI-AML3 and IMS-M2 cells, respectively, and with a higher EC_50_ of ∼ 45 ± 10 μM in HEL cells (**Figure 6D**).

Based on these results, we performed further experiments using the 10 μM dose of EN7 to maximize loss of glutamylation and minimize general cytotoxic effects. We observed evidence of myeloid differentiation in both NPM1c-mutated cell lines as early as day 3 of 10 μM EN7 treatment, which was more pronounced in OCI-AML3 cells by day 6 (**Figure 6E,F**). EN7-treated IMS-M2 and OCI-AML3 cells demonstrated increased expression of the myeloid markers CD13, CD14, and CD11b compared to DMSO controls (**Figure 6E, Supplementary Figure S6F**) and displayed myeloid morphology not observed in untreated cells at the same timepoints (**Figure 6F, Supplementary Figure S6G**). Due to the strong inhibitory effect of EN7 on IMS-M2 proliferation—very few viable cells remained by day 6—we were unable to analyze differentiation in those cells at that time point. Notably, EN7 treatment did not appear to induce myeloid differentiation in NPM1-wildtype HEL cells; expression of myeloid cell surface markers was unchanged relative to controls at both day 3 and day 8, and there were no observable morphological differences between EN7-treated cells and DMSO controls at either timepoint (**Figure 6E,F**). All these data support our hypothesis that the phenotype caused by EN7 is driven by TTLL4 inhibition.

## Discussion

Our study identifies TTLL4-mediated glutamylation as a critical post-translational modification (PTM) regulating NPM1c-dependent leukemogenesis. We found that the NPM1c intrinsically disordered region, specifically the second acidic stretch that is required for binding to chromatin, is hyper-glutamylated by the TTLL4 glutamyltransferase. Loss of TTLL4 through multiple approaches resulted in loss of NPM1c mono-glutamylation and impaired proliferation, while promoting myeloid differentiation of NPM1-mutated leukemic cells. This is accompanied by the upregulation of myeloid gene expression programs in TTLL4-deficient leukemic cells. Mechanistically, TTLL4 loss leads to NPM1/NPM1c relocalization from the cytoplasm to chromatin, which may contribute to the observed differentiation phenotype in NPM1-mutant acute myeloid leukemia (AML). Importantly, we found that genetic deletion of *Ttll4* also promotes myeloid differentiation of leukemic cells *in vivo* in a murine model of NPM1c AML, which is associated with significantly prolonged survival. We do not observe the same dependency on TTLL4 in wildtype hematopoietic progenitors or in NPM1-wildtype leukemic cells, suggesting some selectivity of *TTLL4* inactivation in NPM1c-mutated cells.

Our data is consistent with the hypothesis that loss of glutamylation on NPM1c as a result of *TTLL4* inactivation leads to a re-distribution of NPM1/NPM1c from the cytoplasm to the nucleus, which leads to loss of self-renewal and a gain of myeloid differentiation programs in leukemic cells. NPM1c has been shown to be recruited to chromatin at self-renewal loci by the nuclear export factor XPO1, which binds to the NES generated by the NPM1c mutation^32^. XPO1 is also known to be required for the cytoplasmic localization of NPM1c, which is known to contribute to its transforming capabilities^2,7,21^. Disordered regions like the NPM1c IDR (the site of TTLL4-mediated glutamylation) provide accessibility that facilitates NES binding to XPO1^33^. As *TTLL4* knockdown phenocopies selinexor-induced NPM1c nuclear retention and myeloid differentiation, glutamylation may play a role in XPO1-mediated chromatin interactions and nuclear-cytoplasmic shuttling. It is likely that glutamylation of NPM1c stabilizes its cytoplasmic retention, although the precise molecular mechanism remains unclear and will be an important area of future study.

Further study is necessary to elucidate how exactly TTLL4 glutamylation of NPM1c inhibits differentiation in AML, and whether TTLL4 plays additional roles beyond affecting NPM1c intracellular localization. The A2 acidic stretch, which we observed is uniquely hyper-glutamylated in the mutant protein, has been found to mediate NPM1c binding to *HOX* genes and its interaction with transcription-regulating machinery at those loci to promote gene expression^8,12^. However, we do not observe direct changes in *HOX* or other self-renewal gene expression upon *TTLL4* knockdown. This suggests that TTLL4 influences leukemic gene expression through mechanisms that may be distinct from canonical NPM1c-mediated *HOX* gene regulation. Given the critical role of histone chaperones in chromatin regulation, it is possible that TTLL4-mediated glutamylation modulates NPM1c function beyond direct chromatin interactions and instead influences the histone chaperone activity of NPM1c. Notably, TTLL4-mediated glutamylation of the NPM1 embryonic paralog Npm2 enhances its affinity for histones, raising the possibility that hyper-glutamylation of NPM1c could similarly regulate histone interactions, potentially sequestering histones from self-renewal loci and altering chromatin accessibility in AML^17^. Future studies examining changes in NPM1c chromatin occupancy as well as proteomic analysis of NPM1c interactors in *TTLL4*-deleted cells will be necessary to delineate these functional distinctions and provide further clarification on contribution of glutamylation to aberrant gene expression in AML.

We found that TTLL4 loss does not fully phenocopy NPM1c loss at the gene expression level, suggesting that there may be additional NPM1c-independent functions of TTLL4 contributing to leukemogenesis. Beyond NPM1c, other known targets of TTLL4 have been found to be dysregulated in AML, including the histone chaperone NAP1 and the transcription factor KLF4, a target of PU.1 that contributes to monocytic differentiation^14,17,34-37^. Another TTLL4 substrate is β-tubulin, which plays key roles in mitosis^24,25^. However, we found that TTLL4 loss does not result in gene expression changes related to cell division and does not affect the proliferation of wildtype HSPCs. Consistent with prior observations that *Ttll4* germline deletion is not lethal in mice, this suggests that TTLL4 may be dispensable for normal cellular function and healthy hematopoiesis^14,38,39^. While our findings establish a crucial role for TTLL4 in NPM1c-driven leukemogenesis, several questions remain. The broader impact of TTLL4-mediated glutamylation on global cellular function remains unexplored, and potential compensation by other TTLL family members should be investigated. Additionally, our study highlights the need for rescue experiments restoring TTLL4 activity in knockdown cells to definitively confirm its causative role in leukemic transformation.

Importantly, our work shows that TTLL4 can be targeted with small molecule inhibitors. We report EN7 as a first-in-class small molecule inhibitor of TTLL4 that phenocopies *TTLL4* deletion, significantly inhibiting cellular proliferation and inducing myeloid differentiation in NPM1c-mutant AML cells. Notably, this inhibitor exhibited limited effects on NPM1-wildtype cells, suggesting that TTLL4-mediated glutamylation is a critical regulator of NPM1c-specific mechanisms of leukemogenesis. Given the deep pocket and unique TTLL4 residues, future studies will pursue improved lead inhibitor candidates.

In summary, NPM1 glutamylation and the implication of TTLL4 in chromatin remodeling suggests that histone chaperone glutamylation may contribute to the dysregulation of normal hematopoiesis. Our work demonstrates that TTLL4 preferentially glutamylates NPM1c and identifies TTLL4 as a key regulator of aberrant self-renewal and transformation of NPM1c-mutated leukemic cells. By linking post-translational glutamate-glutamylation to leukemic transcriptional programs, we provide novel insights into the epigenetic regulation of AML pathogenesis. Overall, we conclude that TTLL4 is a promising therapeutic target in NPM1c AML. Future work investigating the therapeutic potential of targeting TTLL4-mediated glutamylation could offer new avenues for differentiation-based therapies in NPM1c-driven leukemia.

## Methods

### Cell lines

All human AML cell lines were cultured in RPMI-1640 with L-glutamine (Corning) supplemented with 10% FBS (Gibco) and 1% penicillin/streptomycin (Corning) at 37°C and 5% CO_2_. OCI-AML3 (RRID: CVCL_1844) with constitutive spCas9 expression (LentiV_Cas9_puro, RRID:Addgene_108100) was a gift from Dr. Roger Belizaire at Dana-Farber Cancer Institute and were cultured with 1 μg/mL puromycin (Cayman) to maintain spCas9 vector expression until lentiviral transduction. NPM1c-FKBP12^F36V-^GFP OCI-AML3 were a gift from Dr. Margaret Goodell^21^. IMS-M2 cells were a gift from Dr. Yogen Saunthararajah. HEK 293T cells (RRID:CVCL_0063) were cultured in DMEM with 4.5 g/L glucose and without L-glutamine and sodium pyruvate (Corning) supplemented with 10% FBS and 1% penicillin/streptomycin. All cell lines were authenticated by STR profiling (Einstein Genomics Core) and routinely tested for mycoplasma by PCR using primers for GPO (5’-ACTCCTACGG-GAGGCAGCAGT-3’) and MGSO (5’-TGCAC-CATCTGTCACTCTGTTAACCTC-3’).

### Experimental animals

*Npm1*^*frt-cA/+*^*;R26*^*FlpoER*^ mice were generated as previously described (The Jackson Laboratory, Strains No. 033164 and No.019016)^29^. For FLPo recombinase induction and expression of the *Npm1*^*cA*^ mutant allele, mice received 125 mg/kg tamoxifen by oral gavage once daily for three consecutive days. Wildtype (C57Bl/6) mice were purchased from Taconic. CD45.1^+^ recipient mice (B6.SJL) were purchased from The Jackson Laboratory (Strain No. 002014). Experimental mice included both males and females. Peripheral blood was obtained under isoflurane anesthesia by retro-orbital bleeding. Complete blood counts were analyzed on Genesis (Oxford Science) blood analyzer. All animal experiments were conducted with the approval of the Institutional Animal Care and Use Committee at Albert Einstein College of Medicine (AECOM; #00001181 and #00001165).

### Covalent crosslinking of total NPM1 antibody to Protein A agarose

Protein A agarose beads (Millipore) were washed with 1X PBS, followed by incubation with total NPM1/NPM1c antibody (ProteinTech) in 1X PBS. The mixture was gently rotated at room temperature for 1 hour, then centrifuged at 300 x *g* for 1 minute, and the supernatant was removed. A 1% BSA solution was added to the antibody-bead conjugate, rotated at room temperature for 1 hour, and centrifuged again to remove the supernatant. The beads were washed four times with 1X PBS before incubation with freshly prepared 25 mg/mL dimethyl pimelidate (DMP) in 0.2 M triethanolamine (TEA) (pH 8.0–8.2 at 22°C) for 30 minutes at room temperature with gentle rotation. After centrifugation and removal of the supernatant, a second round of fresh DMP incubation was performed under the same conditions. The DMP solution was removed, and the antibody-bead conjugate was washed with 1X PBS. Fresh 50 mM ethanolamine in 1X PBS (pH 8 at 22°C) was added to quench the reaction, followed by gentle rotation at room temperature for 30 minutes. The supernatant was discarded, and the beads were washed 5–6 times with 1X PBS before storage in 1X PBS at 4°C. Just before use, the NPM1-Protein A agarose conjugate was briefly treated by removing PBS, washing with 100 mM glycine (pH 2.5) and 100 mM Tris (pH 8), followed by three quick washes with 1X PBS. Gamma globulin (Invitrogen) was similarly crosslinked to Protein A agarose bead for use as a negative control.

### Immunoprecipitation

OCI-AML2, OCI-AML3, and IMS-M2 cells were harvested, washed twice with ice cold PBS, and resuspended in 1X RIPA buffer containing 1% NP-40, 150 mM NaCl, 1 mM EDTA, 50 mM Tris-HCl (pH 8, 4°C), 0.25% sodium deoxycholate, 0.1% SDS, along with protease, phosphatase, and acetylase inhibitors (ThermoFisher). Lysates were incubated on gentle rotation at 4°C for 30 minutes with intermittent vortexing, followed by centrifugation to remove cellular debris. The supernatant was carefully collected into a new low-adhesion tube, avoiding disturbance of the pellet and pre-cleared by incubating with Protein A agarose bead slurry (Pierce) for 1h at 4°C with gentle mixing. The pre-cleared supernatant was collected by centrifugation at 10,000 x *g* for 10 minutes and the concentrations were normalized by bicinchoninic acid assay (Pierce) to 3-5 mg per condition (2 μg/μL). Total NPM1-Protein A agarose bead conjugate was added to the supernatant and incubated overnight at 4°C with gentle mixing. Gamma globulin-Protein A agarose bead conjugate was used as a negative control. The bound protein was subjected to low pH glycine elution. Briefly, the beads were centrifuged at 300 x *g* at 4°C and washed sequentially with 1X PBS, four times with 1X PBS + 0.05% Tween-20, and once with Milli-Q water. For glycine elution, 0.1 M glycine (pH 2.5, 200–300 μL) was added and incubated for 15 minutes at room temperature with gentle mixing. Beads were centrifuged at 300 x *g* at room temperature, and the supernatant, containing glycine-eluted NPM1/NPM1c protein, was immediately neutralized with 1 M Tris (pH 8) and stored at -20°C for long-term use. This glycine elution was repeated three times in total. Samples were prepared in 1x Laemmli buffer for immunoblotting with antibodies as specified.

### Immunoblotting

Human cells were washed once with room temperature 1X PBS and lysed using RIPA buffer (1% NP-40, 150 mM NaCl, 1 mM EDTA, 50 mM Tris-HCl (pH 8, 4°C), 0.25% sodium deoxycholate, 0.1% SDS) along with 1X protease inhibitor cocktail. The samples were mixed with 1X Laemmli Buffer and denatured for 15 minutes at 95°C. Samples were loaded onto Any kD Mini-Protean TGX gels (Bio-Rad) with a pre-stained protein ladder (ThermoFisher). Proteins were separated by electrophoresis for 40 minutes and transferred onto PVDF membranes (Immobilon, Millipore), followed by blocking using a 1% ECL Prime blocking agent (Cytiva). The blots were incubated overnight at 4°C with total NPM1/NPM1c (ProteinTech), NPM1c (Novus Biologicals), TTLL4 (Invitrogen), TTβIII/α-mono-glutamylation, β-actin (Sigma-Aldrich), or vinculin (Sigma-Aldrich) antibody. Membranes were next washed in PBS-T (1X PBS + 0.01% Tween-20) and incubated with anti-rabbit or anti-mouse IgG HRP secondary antibody (Cytiva) followed by detection using ECL chemiluminescence (Lumigen, TMA-6) and imaging with an ImageQuant LAS4000 system (GE). Mouse cells were washed once with room temperature 1X PBS and lysed for 30 minutes on ice using NP-40 Cell Lysis Buffer (1% NP-40, 50 mM Tris-HCl (pH 7.4), 150 mM NaCl, 5 mM EDTA) (Alfa Aesar) supplemented with 0.1 mM Na3VO4, 1 mM PMSF, and 1X Pierce protease and phosphatase inhibitor (ThermoFisher). Lysates were run on NuPAGE 10% Bis-Tris protein gels (ThermoFisher) at 200 V with NuPAGE 1X MOPS SDS Running Buffer (ThermoFisher) and transferred onto a nitrocellulose membrane at 30V with NuPAGE 1X Transfer Buffer (ThermoFisher) supplemented with methanol and NuPage antioxidant (ThermoFisher). Membranes were blocked with 5% TBS-Tween 20 (TBS-T) in blotto non-fat milk for 1 hour at room temperature. Membranes were then incubated with rabbit primary antibodies against TTLL4 (Invitrogen), TTβIII/α-mono-glutamylation, total NPM1/NPM1c (ProteinTech), or NPM1c (Novus Biologicals) in 5% TBS-T in BSA overnight at 4°C. This was followed by incubation with goat loading control primary antibodies against vinculin and β-actin in 5% TBS-T in BSA for 1 hour at room temperature followed by incubation with anti-rabbit IgG HRP (Cell Signaling Technologies) and anti-goat IgG IRDye 680RD (LiCor) secondary antibodies in in 5% TBS-T in BSA for 1 hour at room temperature. Membranes were briefly washed in ECL solution containing 250 mM luminol, 90 mM p-coumaric acid, 1 M tris (pH 8.5), and 30% hydrogen peroxide before imaging using LI-COR software.

### Generation of TTβIII/α-mono-glutamylation antibody

A new rabbit polyclonal antibody specifically recognizing mono-glutamylated epitopes was generated according to method previously described by Spano and Frankfurter^40^. Briefly, rabbits were immunized with the synthetic peptide antigen MY{Glu(Glu)}DDEEESEAQGPKC, corresponding to the mono-glutamylated sequence, conjugated to the carrier protein keyhole limpet hemocyanin (KLH). The initial immunization regimen consisted of three rounds of immunization. To improve antibody titer, an additional four rounds of boosting immunizations were performed. Following immunizations, antisera were affinity-purified against the glutamylated peptide antigen and cross-adsorbed using the non-glutamylated control peptide MYEDDEEESEAQGPKC to remove antibodies recognizing unmodified epitopes. Antibody production and purification were performed by GenScript USA Inc. (Piscataway, NJ). Validation of antibody specificity for glutamylation is shown in Supplementary Figure S7 using Nap1 and NPM2 proteins treated with or without TTLL4.

### Expression and purification of recombinant proteins

N-terminally GST fused recombinant TTLL4 catalytic domain (561-1199aa) was expressed and purified as described earlier^17^. Nap1L4, NPM1 and NPM1c full-length protein were expressed in BL21 (DE3) competent *E. coli* cells grown in LB medium containing appropriate antibiotics (GoldBio). Cultures were grown to OD_600_ ∼0.7 at 37°C with agitation before induction with 0.5 mM IPTG (GoldBio) followed by a 16-hour incubation at 20°C with agitation. Cells were harvested by centrifugation and resuspended in lysis buffer (20 mM Tris–HCl pH 8, 300 mM NaCl, 5% glycerol, and 5 mM β-mercaptoethanol). PMSF (0.2 mM), 1X protease inhibitor cocktail (Roche), and 1 μg/mL DNase were added to cell suspension and the cells were lysed using an Emulsiflex C3 homogenizer (Avestin) at 4°C with two passages at 12,000 psi. The lysates were centrifuged at 14,000 × *g* for 45 minutes and the cleared lysates were loaded onto HisPur Ni-NTA (Thermo) gravity-flow column. The columns were washed extensively with 20 column volumes (CVs) of lysis buffer, followed by 5 CVs of the same buffer containing 30 mM and 45 mM imidazole; proteins were eluted with 2 CVs of the same buffer containing 300 mM imidazole. Eluate containing Nap1L4, Npm1 and Npm1c were transferred to dialysis tubing (10,000 MWCO) and the N-terminal His_6_ affinity tag was cleaved by adding 6 ×-histidine-tobacco etch virus (TEV) protease catalytic domain at a 1:50 mass ratio (TEV:NPM1) and dialyzed for 16 hours at 4°C against buffer containing 50 mM Tris-HCl pH 8.0, 150 mM NaCl, 2 mM DTT, and 10% glycerol. Subtractive Ni-NTA chromatography was then performed to remove the affinity tag and TEV protease. Proteins were concentrated to ∼10 mg/mL and subjected to size exclusion chromatography (Hiload^TM^ 16/600 Superdex^TM^ 200 pg, GE) using the aforementioned buffer. The NPM1c protein was subjected to an additional anion exchange chromatography on MonoQ column (Cytiva). Fractions containing pure protein were pooled and concentrated to ∼10 mg/mL and stored at −80°C. Protein concentrations were measured by Bradford protein assay reagent using BSA as standard.

### In vitro glutamylation of NPM1-wildtype and NPM1c-mutant protein

The reaction was carried out as previously described^17^. Briefly, NPM1 (wildtype or mutant), and recombinant human TTLL4, at a mass ratio of 1:100 (TTLL4:NPM1) were incubated in the assay buffer containing 50 mM Tris, pH 8.5, 4 mM MgCl2, 100 mM NaCl, 10 mM β-mercaptoethanol, 5% glycerol, 2.5 mM K^+^-L-Glutamate, and 2.5 mM ATP for 2 hours at 30°C. At the end of 2 hours, another aliquot of 1:100 TTLL4 was added, and the mixture was incubated for an additional 2 hours. Glutamylation was assessed via immunoblot. TTLL4 was removed from the reaction mixture by passing through a Ni^2+^-NTA column. Glutamylated NPM1 proteins were dialyzed extensively against 20 mM Tris-HCl buffer, pH 8, containing 100 mM NaCl, 2 mM DTT, 1 mM EDTA, 1X protease inhibitor cocktail and stored at −80°C.

### Mass spectrometry sample preparation

Recombinant proteins (glutamylated NPM1 and NPM1c) and IMS-MS2 (cellular NPM1/NPM1c) samples were subjected to in-tube trypsin digestion by diluting samples 10-fold in 100 mM ammonium bicarbonate (pH ∼8), reducing disulfide bonds with 25 mM DTT at 50°C for 30 minutes, and alkylating cysteines with 75 mM iodoacetamide in the dark for 1 hour. Proteins were digested with trypsin (1:100 enzyme-protein mass ratio) at 37°C for 4 hours, then frozen until use. OCI-AML2 (cellular NPM1) and OCI-AML3 (cellular NPM1/NPM1c) samples were vacuum concentrated and subjected to in-column trypsin digestion by diluting 5-fold in a buffer containing 5% SDS, 5 mM DTT and 50 mM ammonium bicarbonate (pH ∼8) and incubated 1 hour at room temperature for disulfide bond reduction. Samples were alkylated with 20 mM iodoacetamide in the dark for 30 minutes, followed by addition of aqueous phosphoric acid to a final concentration of 1.2%. Samples were then diluted in six volumes of binding buffer (90% methanol and 10 mM ammonium bicarbonate, pH 8.0). After gentle mixing, the protein solution was loaded to an S-trap filter (Protifi) and spun at 500 g for 30 seconds. The sample was washed twice with binding buffer. Finally, 1 μg of sequencing grade trypsin (Promega), diluted in 50 mM ammonium bicarbonate, was added into the S-trap filter and samples were digested at 37°C for 18 hours. Peptides were eluted in three steps: (i) 40 μL of 50 mM ammonium bicarbonate, (ii) 40 μL of 0.1% TFA, and (iii) 40 μL of 60% acetonitrile and 0.1% TFA. The peptide solution was pooled, centrifuged at 1,000 x *g* for 30 seconds and dried in a vacuum centrifuge.

### LC-MS/MS acquisition and analysis

Hydrophilic interaction liquid chromatography (HILIC) was used to clean up cellular samples for mass spectrometry analysis as described previously, cleaned by hydrophilic interaction liquid chromatography (HILIC) with additional ammonium formate in 90% acetonitrile rinses to remove polyethylene glycol^41^. Samples were reconstituted in 20 μL HPLC solvent A (0.3% formic acid) for LC-MS analysis using custom-packed analytical and pre-columns (Agilent). A 1100 Series Binary HPLC system (Agilent Technologies) coupled to an Orbitrap Fusion Tribrid mass spectrometer (ThermoFisher) was used for LC-MS analysis of the samples^42^. Reverse-phase separation was performed with a gradient from 0%-60%-100% Solvent B (0.3% formic acid in 72% acetonitrile, 18% isopropanol) over 65 minutes at ∼100 nL/min. Pre-columns were pressure loaded with 100 fmol each of Angiotensin I and Vasoactive Intestinal peptide standards and with either 10% of the ISM-MS2 digested sample, 25% AML2 or AML3 digested sample, or 5% of recombinant NPM1 or NPM1c glutamylated sample. A full mass spectrum was acquired in the Orbitrap with 60,000 resolution and mass range of 300-2000 m/z. Precursor ions were selected for fragmentation using a decision tree based on charge state. Briefly, CAD and ETD for charge state 2-3 ions (ion trap analysis)^43^, stepped HCD (27%) and ETD for charge 4 and higher ions (Orbitrap analysis), with dynamic exclusion for 12 seconds. Intact analysis of recombinant NPM1/NPM1c was conducted using a modified Orbitrap Velos Pro^TM^ upgraded to an Orbitrap Elite, incorporating proton transfer charge reduction (PTCR) with perfluoro(methyldacalin) for enhanced analysis^44^. Proteins (∼100 fmol) were pressure-loaded and eluted for over 25 minutes, with PTCR targeting 800 m/z and a 400 Da isolation range (3000-3500 m/z). Byonic (Protein Metrics) was used for MS data analysis against NPM1/NPM1c isoforms and a contaminants database, incorporating semi-specific cleavage and modifications such as single and double glutamylation of glutamic acid, phosphorylation of serine, threonine, and tyrosine, mercaptoethanol- or carbamidomethyl-addition to cysteine, and oxidation of methionine. Peptide-spectra matches guided manual verification of glutamylation sites.

### CRISPR-Cas9 TTLL4 deletion in OCI-AML3 cells

Human *TTLL4* sgRNA and non-targeting (sgNT) control oligos were designed using the Broad Institute’s “CRISPick” software^45,46^ and cloned them into the pLKO.5.sgRNA.EFS-tRFP (RRID:Addgene_57823) lenti-viral vector through the Azenta CRISPR Construct Synthesis service (sgTTLL4-1: 5’-TAAGTTATTCCGGTTATACC-3’; sgTTLL4-2: 5’-ACCG-GAATAACTTAGCCATG-3’; sgNT: 5’-GCACTAC-CAGAGCTAACTCA-3’). Lentiviral particles containing the sgTTLL4 or sgNT expression vectors were produced in HEK293T cells using pCMVR8.74 packaging plasmid (RRID:Addgene_22036) and pCMV-VSVg empty backbone plasmid (RRID:Addgene_8453) with the 293Tran transfection reagent (OriGene) according to the 293Tran manufacturer protocol. Cas9-OCI-AML3 cells were transduced by incubation with lentiviral supernatant and 8 μg/mL polybrene transfection reagent (MilliporeSigma). RFP^+^ cells were isolated 3 days post-transduction using fluorescent-activated cell sorting (FACS); this time-point marks day 0 for all experiments using these cells. *TTLL4* editing was validated six days after FACS by Azenta AmpliconEZ next-generation sequencing. DNA was isolated from cells using the DNeasy Blood Mini Kit (Qiagen) and human *TTLL4* exon 3 was amplified using primers surrounding the sgRNA cut sites (hTTLL4-ex3_FW: 5’-CAGTGCCCATATCGCCTTGT-3’, hTTLL4-ex3_RV: 5’-GAGGCAAGGAGGTGAATGCT-3’). PCR products were purified using the Monarch^®^ PCR & DNA Cleanup Kit (NEB) before sending to Azenta for sequencing. Loss of NPM1c glutamylation was validated six days after FACS by immunoblot analysis with TTβIII/α-mono-glutamylation antibody.

#### shRNA TTLL4 knockdown in NPM1c-degron OCI-AML3 cells

*TTLL4* shRNA oligo pairs (5’-CCGGGTGGCCAC-GCAGCCTTATAAACTCGAGTTTATAAGGCTGCG TGGCCACTTTTT-3’; 5’-AATTAAAAAGTGGCCAC-GCAGCCTTATAAACTCGAG-TTTATAAGGCTGCGTGGCCAC-3’) were reconstituted to 0.1 nmol/μL in annealing buffer (100 mM NaCl, 10 mM Tris-HCl pH 7.4), brought to a boil, then slowly cooled to room temperature followed by 1:400 dilution in 0.5x annealing buffer. The pLKO-Tet-On vector (RRID:Addgene_21915) was digested using AgeI and EcoRI then gel purified (NEB). Ligation was performed by mixing 0.01 pmol/μL vector and 0.1 pmol/μL insert with Quick T4 Ligase (NEB) in T4 Ligase Buffer (500 mM Tris pH 7.5 at 25°C, 100 mM MgCl2, 10 mM ATP, 100 mM DTT) followed by transformation into *E. coli*. Constructs were verified with Sanger sequencing. Lentiviral particles containing the pLKO-Tet-On-shTTLL4 or scrambled control (shScr) expression vectors were produced in HEK293T cells using calcium phosphate transfection followed by ultracentrifugation to concentrate the viral supernatant. NPM1c-FKBP12^F36V-^GFP OCI-AML3 cells were transduced by centrifugation with lentivirus at 30°C, 500 x *g* for 90 minutes with 2 μg/mL polybrene followed by selection with 2 μg/mL puromycin (Cayman) starting at 48 hours. Successful *TTLL4* knockdown was confirmed by *TTLL4* RT-qPCR (shTTLL4-1_FW: 5’-TCTCTCCCG-GACTTGTTCAAC-3’, shTTLL4-1_RV: 5’CGAACGCAA-GCAGAAAGACTC-3’; shTTLL4-2_FW: 5’-AGACTCAAGCTGGCCTTTCC-3’, shTTLL4-2_RV: 5’-CAGTTATGGGCTCACAGCCA-3’) and by immunoblot analysis with TTβIII/α-mono-glutamylation antibody and NPM1/NPM1c antibody (ProteinTech) as a loading control.

### CRISPR-dCas9 TTLL4 knockdown in OCI-AML3 cells

A two-plasmid lentiviral system was employed for the knockdown of the target gene *TTLL4*. The lenti_dCas9-KRAB-MeCP2 plasmid was a gift from Andrea Califano (RRID: Addgene_122205)^47^, while pXPR_050 was a kind gift from John Doench & David Root (RRID: Addgene_96925)^45^. sgRNA sequences were designed using the Broad Institute’s “CRISPick” software^45,46^ and cloned into pXPR_050 following published protocol (sgTTLL4-1: 5’-ACCGAG AGTCTGGAAGAAGA-3’, sgTTLL4-2: 5’-CACAC-CGCGCGCGCCCCCAG-3’, sgNT: 3’-GTGTAG-TTCGACCATT CGTG-3’)^45^. Lentiviral particles were generated and transduced into OCI-AML3 cells as previously described by our group^48^. Stable expression of the dCas9-KRAB-MeCP2 construct in OCI-AML3 cells was maintained through blasticidin selection (10 μg/mL; Cayman). Subsequently, the cells were transduced with the sgRNA constructs followed by puromycin selection (2 μg/mL; Cayman). The timepoint at two days post selection marks day 0 for all experiments using these cells. Successful *TTLL4* knockdown was confirmed by *TTLL4* RT-qPCR (sgTTLL4-FW: 3’-AGACTCAAGCTGGCCTTTCC-5’, sgTTLL4-RV: 3’-CAGTTATGGGCTCACAGCCA-5’) and by immunoblot analysis with TTβIII/α-mono-glutamylation antibody using total NPM1/NPM1c (ProteinTech) and vinculin (Sigma-Aldrich) antibodies as loading controls.

### RT-qPCR validation of TTLL4 knockdown

*TTLL4* knockdown cells were collected after selection and RNA was extracted using the Monarch Total RNA Kit (NEB) following the manufacturer’s protocol. 1 or 2 μg of RNA was used as an input for cDNA synthesis using the M-MLV reverse transcriptase kit (Invitrogen). The synthesized cDNA was used for real-time quantification of *TTLL4* transcript levels using the 2X Universal SYBR green fast qPCR mix (ABclonal). The standard comparative Ct with melt program on ThermoQuant Studio 6 Pro was run. Actin was used as housekeeping control (actin_FW: 5’-AGCTA CGAGCTGCCTGAC-3’, actin_RV: 5’-AAGGTAG-TTTCGTGGATGC-3’). All experiments were conducted in triplicates.

### RNA isolation and sequencing of shRNA TTLL4 knock-down cells

Doxycycline-inducible shTTLL4-2 or control shScr cells were cultured as above. Cells were treated with 250 nM doxycycline hyclate (Cayman) with or without 10 nM dTAG-13 (Tocris) for a total of 96 hours; doxycycline was replenished after 48 hours. RNA was extracted using RNeasy Mini Kit (QIAGEN) following the manufacturer’s protocol. RNA quantification and quality control were accomplished using the Bioanalyzer 2100 (Agilent Technologies). Unstranded RNA-seq libraries were created by Novogene. Barcoded libraries were sequenced on an Illumina platform using 150 nt paired-end libraries generating approximately 30 million paired-end reads (9GB total data) per biological replicate. Reads were trimmed and aligned to the human genome (hg38) using STAR^49^ in the nf-core/rnaseq pipeline (v3.14)^50^. Differential abundance analysis was performed using salmon^51^ read counts and DESeq2^52^ with a custom script (https://github.com/Shechterlab/nextflow-rnaseq). Overenrichment analysis (ORA) and gene set enrichment analysis (GSEA) were performed using ClusterProfiler^53^ and msigdb^54^ in R version 4.2.2 with custom scripts (https://github.com/Shechterlab/Schurer_Ilyas_etal_2025/). Comparisons using GeneOverlap were made between TTLL4*kd*, NPM1c dTAG treatment, and GSE181176 (selinexor, an XPO1 inhibitor), GSE144759 (MI503, a menin inhibitor), GSE228307 (ziftomenib, a menin inhibitor), GSE185091 (TDI055, an ENL inhibitor), GSE205802 (WM1119, a KAT6a inhibitor), GSE132244 (CPI203, a BRD4 inhibitor).

### Sub-cellular fractionation of human AML cell lines

Fractionation was performed as previously described^55^. Briefly, equal number of cells were harvested and washed with PBS. All subsequent steps were performed at 4°C using freshly prepared buffers. Cells were suspended in 300 μL of Hypotonic Lysis Buffer (10 mM Tris-HCl pH 8 at 4°C, 0.1% NP-40, 1 mM KCl, 1.5 mM MgCl_2_, 0.5 mM EDTA, 0.5 mM DTT, and 1X protease inhibitor cocktail) and rotated for 15 minutes at 4°C to facilitate lysis. The lysate was then overlaid onto a 24% sucrose cushion, a density gradient solution consisting of 10 mM Tris, 15 mM KCl, 1 mM EDTA, 24% Sucrose (w/v), 0.5 mM DTT, and 1X protease inhibitor cocktail, and centrifuged at 10,000 x *g* for 10 minutes. The cytoplasmic fraction was carefully collected from the top of the gradient and the nuclei pellet was washed in a PBS/1mM EDTA solution by centrifuging at 10,000 x *g* for 5 minutes. Nuclei pellets were then resuspended in glycerol buffer (20 mM Tris, 75 mM NaCl, 0.5 mM EDTA, 50% Glycerol (v/v), 0.85 mM DTT, and 1X protease inhibitor) and subsequently lysed using equal volume of nuclei lysis buffer, which contained 20 mM HEPES, 1 mM DTT, 7.5 mM MgCl_2_, 0.2 mM EDTA, 300 mM NaCl, 1% NP40, 1M Urea, and 1X protease inhibitor cocktail. Chromatin was spun down by centrifugation at 10,000 x *g* for 5 minutes and the supernatant was collected for the nucleoplasm fraction. Subsequently, the chromatin fraction pellet was washed with PBS/1 mM EDTA and resuspended in RIPA lysis buffer supplemented with protease inhibitor cocktail using probe-tip sonication at 20% amplitude for 5 seconds. Fractions were then analyzed by immunoblotting as described above. Antibodies used for loading controls were β-actin (Sigma-Aldrich) and vinculin (Sigma-Aldrich) for the cytoplasmic fraction, U1-70K (Santa Cruz Biotechnology) for the nucleoplasm fraction, and histone H3 for the chromatin fraction (Abcam).

### Isolation of mouse bone marrow for CRISPR-Cas9

*Npm1*^*cA/+*^, wildtype, or *Npm1*^*cA/+*^;*Nras*^*G12D/+*^ mouse bone marrow was harvested from donor mice and subjected to red blood cell lysis (Qiagen). *Npm1*^*cA/+*^ or wildtype c-Kit^+^ bone marrow cells were enriched from whole bone marrow using CD117 microbeads and MACS magnetic separation (Miltenyi Biotech) per manufacturer’s instructions.

### Generation of NPM1c;NRAS^G12D^ leukemic mice

A total of 300,000 *Npm1*^*cA/+*^;*Nras*^*G12D/+*^ frozen bone marrow cells (provided as a gift from Dr. Linde Miles) and 700,000 CD45.1^+^ B6.SJL helper bone marrow cells were transplanted into 6-to 8-week old lethally-irradiated (9 Gy) B6.SJL recipient mice for expansion of viable leukemic cells. Donor bone marrow was retro-orbitally injected and recipient mice were given 100 mg/mL Baytril-100 (Bayer) in drinking water for 4 weeks after transplantation. Mice were euthanized upon development of AML and whole bone marrow was harvested for CRISPR-Cas9 *TTLL4* deletion.

### CRISPR-Cas9 Ttll4 deletion in mouse bone marrow

Mouse *Ttll4* sgRNA multi-guide oligos and non-targeting sgNT oligo were designed by Synthego (sgTTLL4: 5’-UUCAAGCAGAGACAUCCCUC-3’, 5’-GACCAGACU-UUGACCUCCGA-3’, 5’-AC-GUAGGCACACUGUCAGCA-3’; sgNT: 5’-GCACUACCAGAGCUAACUCA-3’). 35 μM sgTTLL4 or sgNT was incubated with 7.8 μM recombinant spCas9-GFP protein (IDT) for 10-15 minutes at room temperature to form ribonucleoprotein (RNP) complexes. *Npm1*^*cA/+*^ or wildtype c-Kit^+^ bone marrow or *Npm1*^*cA/+*^;*Nras*^*G12D/+*^ whole bone marrow cells were pre-cultured for 1 hour at 37°C in PVA media, comprised of Ham’s F-12 media (Gibco) supplemented with 1% PVA (ThermoFisher), 10 mM HEPES (Life Technologies), 1X ITS-X (ThermoFisher), 2 mM L-glutamine (Gibco), 10 ng/mL mSCF (PeproTech), 10 ng/mL mTPO (Peprotech), and 1% penicillin/streptomycin (Corning). RNPs were electroporated into 1 x10^6^ *Npm1*^*cA/+*^ or wildtype c-Kit^+^ bone marrow or *Npm1*^*cA/+*^;*Nras*^*G12D/+*^ whole bone marrow cells in Buffer T (ThermoFisher) with 30 μM Alt-R electroporation enhancer (IDT) using the Neon NxT transfection system (ThermoFisher) using the following electroporation conditions: 1900 V, 20 ms, 1 pulses^21,56^. After electroporation, cells were plated in PVA media without antibiotics. *Ttll4* editing was validated 24-48 hours after electroporation by ICE analysis^57^ of Sanger sequencing data using primers designed by Synthego. DNA was isolated from cells using the DNeasy Blood Mini Kit (Qiagen) and the region containing *Ttll4* indels was amplified by PCR primers surrounding the sgTTLL4 cut sites (mTTLL4-Synthego_FW: 5’-GCTGGTGCTAGAGGTGTGT-3’; mTTLL4-Sythego_RV: 5’-TTTTGCCTCACGTTGGTGC-3’). PCR products were purified using the Monarch^®^ PCR & DNA Cleanup Kit (NEB) and sent to Azenta for Sanger sequencing using a primer that binds upstream of the sgTTLL4 cut sites (mTTLL4-Synthego_seq: 5’-CTGG TGCTAGAGGTGTGTGG-3’). *Ttll4* editing was also validated 24-48 hours after electroporation by immunoblot using TTLL4 antibody (Invitrogen) and using vinculin antibody (Sigma-Aldrich) as a loading control.

### In vivo bone marrow transplantation of Ttll4-deleted NPM1c;NRAS^G12D^ bone marrow cells

After electroporation of CRISPR-Cas9 RNPs as described above, 250,000 sgNT or sgTTLL4 *Npm1*^*cA/+*^;*Nras*^*G12D/+*^ cells were transplanted into 6-to 8-week old sublethally-irradiated (4.5 Gy) B6.SJL recipient mice for analysis of disease burden and survival.

### CellTiter-Glo^®^ proliferation assays

Cell viability after CRISPR-Cas9 deletion or shRNA knockdown of *TTLL4* was measured using the CellTiter-Glo^®^ Luminescent Cell Viability Assay (Promega). 1 x 10^4^ cells were plated in 100 μL of culture media in 96-well plates and incubated for up to 3 days at 37°C. 100 μL of culture medium without cells was plated to obtain background luminescence. Cells were passaged 1:4 by volume in fresh culture medium every 3 days for set up of additional plates. Plates were equilibrated for 30 minutes at room temperature and then incubated with 100 μL per well of CellTiter-Glo^®^ reagent in the dark for 20 minutes at room temperature with gentle rotation. Luminescence was measured with bidirectional reading and emission at 520 nm on the Victor X plate reader.

### Methylcellulose colony-forming unit (CFU) assays

OCI-AML3 cells, *Npm1*^*cA/+*^ or wildtype c-Kit^+^ bone marrow cells were resuspended in IMDM with L-glutamine and 25 mmol/L HEPES media (Life Technologies) supplemented with 10% FBS and 1% penicillin/streptomycin. For OCI-AML3 cells, 2.5 x10^4^ cells in IMDM were added to 5 mL MethoCult H4435 enriched methylcellulose media (StemCell) and plated at a final concentration of 5 x10^3^ cells per dish. For mouse c-Kit^+^ bone marrow cells, 1.5 x10^4^ cells in IMDM were added to 5 mL mouse HSC007 methylcellulose complete media (R&D) and plated at a final concentration of 3 x10^3^ cells per dish. Plates were incubated at 37°C and CFUs were counted 7 days post-plating.

### Flow cytometry analysis of human cell lines

For analysis of myeloid differentiation, cells were stained with anti-human Fc block antibody (clone Fc1, BD Biosciences) at 4°C for 10 minutes. Cells were then stained with anti-human CD11b (clone ICRF44 or M1/70, BioLegend), CD13 (clone WM15, BioLegend), and CD14 (clone M5E2, BioLegend) fluorochrome-conjugated surface antibodies at 4°C for 30 minutes. DAPI viability dye (MP Biomedicals) was then added to cells at a 1:10,000 dilution. Flow cytometry was performed on the CYTEK AURORA and analyzed using FlowJo software (V10).

### Flow cytometry analysis of mouse cells

For analysis of immature and mature populations in *Npm1*^*cA/+*^;*Nras*^*G12D/+*^ mice, peripheral blood or bone marrow from crushed bones were subjected to RBC lysis (Qiagen). Lysed peripheral blood cells were stained with anti-mouse CD16/CD32 Fc block antibody (clone 2.4G2, BD Biosciences) at 4°C for 10 minutes then stained with anti-mouse CD117/c-Kit (clone 2B8, BioLegend) and CD45.2 (clone 104, BD Biosciences) fluorochrome-conjugated surface antibodies at 4°C for 30 minutes. Lysed bone marrow cells were stained for 30 minutes at 4°C with lineage (Lin) cocktail of anti-mouse biotin-labeled antibodies: B220 (clone RA3-6B2, Invitrogen), CD19 (clone MB19-1, BioLegend), CD3ε (clone 145-2C11, BioLegend), Gr1 (clone RB6-8C5, BioLegend), and Ter119 (clone TER-119, BioLegend). Lineage-stained cells were then stained for 30 minutes at 4°C with anti-mouse fluorochrome-conjugated surface antibodies: streptavidin (BioLegend), Sca1 (clone D7, BioLegend), CD117 (clone 2B8, BioLegend), and CD45.2 (clone 104, BD Biosciences). For analysis of myeloid differentiation, lysed bone marrow cells were stained for 30 minutes at 4°C with antimouse fluorochrome-conjugated surface antibodies: CD117 (clone 2B8, BioLegend), CD11b (clone M1/70, Invitrogen), CD115 (clone AFS98, BioLegend), F4/80 (clone BM8, BD Biosciences), and CD45.2 (clone 104, BD Biosciences). Flow cytometry was performed on the CYTEK AURORA and analyzed using FlowJo software (V10).

### Histopathology analysis of cytospins and blood smears

To prepare human AML cell line cytospins, approximately 100,000 cells were collected and washed once in PBS before deposition onto a microscope slide using the StatSpin CytoFuge 2 Cytocentrifuge (HemoCue). Cyto-spin preparations of human AML cells and blood smears prepared from peripheral blood of *Npm1*^*cA/+*^;*Nras*^*G12D/+*^ mice were stained with Wright-Giemsa Hema 3 stain (ThermoFisher). Images were captured on a Zeiss Axiovert 200M microscope.

### Analysis of moribund mice

Mice were euthanized upon meeting endpoint criteria of decreased activity, palpable splenomegaly, ruffled fur, and/or body condition score (BCS) <2. Peripheral blood was collected prior to euthanasia and spleen, liver, and bone marrow were harvested post-mortem. Complete blood counts were analyzed on Genesis (Oxford Science) blood analyzer and liver and spleen were weighed for confirmation of AML.

### Structural modeling of TTLL4 catalytic domain

The catalytic domain of human TTLL4 (S533-F1106) was modeled using AlphaFold Multimer, guided by structural homology with mouse TTLL6 crystal structures (PDB: 6VZW, 6VZT)^31,58^. The AlphaFold model was validated by superimposing known TTLL6 structures. The modeled catalytic site included ATP-binding residues (K721, R727-728, K762-764, T809-810), glutamate-binding residues (R727, R788, K924), and magnesium-binding residues (D893, E906, N908). The highest ranked model had the acidic tail of *Xenopus tropicalis* Nap1L4 docked into the predicted catalytic site, aligning its glutamate residue with the active site. ATP and the 2TI intermediate from TTLL6 were overlaid to validate substrate and ligand binding.

### Ligand-based and structure-based virtual screening

We contracted with Cayman Chemical (Ann Arbor, MI) to perform ligand sed and structure-based virtual screening of approximately 9 million compounds commercially available from the Aldrich Market Select^®^ (MilliporeSigma) library using Schrödinger software. For ligand-based screening, the phosphorylated initiation analog (PIA), structurally related to the TTLL6 reaction intermediate 2TI, served as the reference structure. Compounds were initially filtered based on shape similarity to PIA, and the top 2% (200,000 compounds) underwent Glide docking against the TTLL4 catalytic site using a docking grid encompassing the active site residues within a 25 Å box. Following docking, the top 1% of hits were ranked by Molecular Mechanics-Generalized Born Surface Area (MM-GBSA) calculations to estimate binding free energies, refining the list to approximately 650 compounds. Hit molecules were clustered based on chemical similarity, and representative molecules from these clusters were selected based on their predicted binding energy, drug-likeness, and commercial availability, resulting in 56 compounds for biochemical testing. We also contracted with TargetMol Chemicals Inc. (Boston, MA) to conduct a separate structure-based virtual screening. TargetMol screened approximately 1.5 million compounds from the ChemDiv database, alongside the bioactive (L4000) and natural products (L6020) libraries. The TTLL4 catalytic domain model was processed and optimized using Apopdb2receptor tool from OpenEye Scientific Software (Release 3.2.0.2). OEDock software (OpenEye Scientific Software, Ver 3.2.0.2) was then employed to dock compounds into a receptor grid defined around key catalytic residues. Compounds with docking scores below -6 kcal/mol were prioritized, resulting in 104,345 hits. Further filtering based on drug-like properties (Lipinski’s rules, logP, TPSA, CYP metabolism) yielded 676 compounds, which were further refined based on chemical similarity searches against ATP and 2TI analogues, ultimately selecting 114 compounds for biochemical testing. Further confirmation of EN7 and TM42 protein-ligand docking was performed using Schrödinger Glide software. Virtual screen hit compounds were purchased from TargetMol, MilliporeSigma, VitasM, ChemDiv, and Enamine. The Nucleoside Mimetic Library NML-320 (Enamine, Ukraine) was also purchased for screening.

### TTLL4 activity and inhibitor screening

Assayed compounds include the purchased hits from the virtual screen and the 320 compound in the Nucleoside Mimetic Library NML-320 (Enamine, Ukraine). TTLL4 activity and inhibition assays were performed in 50 mM Tris buffer, pH 8.5 containing 50 mM NaCl, 10 mM 2-mercaptoethanol, 4 mM MgCl_2_ at 30°C. For chemical library screening, 1 μg TTLL4 in 45 μL assay buffer was incubated with 100 μM compound for 5 minutes. The glutamylation reaction was initiated by the addition of 5 μL substrate mixture containing 25 μg Nap1L4, 200 μM ATP and 1 mM glutamate. After 10 minutes the reaction was quenched by the addition of 5 μL 0.5 M EDTA. 1 μL of the quenched reaction mixture was spotted onto nitrocellulose membrane. The membrane was washed 3X with PBST, blocked with ECL Prime Blocking Reagent (Cytiva), incubated with TTβIII/α-mono-glutamylation antibody (1:10,000), and detected by ECL chemiluminescence (Lumigen, TMA-6) and imaged with an ImageQuant LAS4000 (GE). IC_50_ determination assays were performed as above except inhibitor concentrations were varied and 2 μM NPM1/NPM1c were used as acceptor substrates. Densitometric quantifications were performed using ImageJ v1.54p with MicroArray_Profile.jar plugin. Further analysis of hit compounds EN7 (from Enamine NML library) and TM42 (from the structure based virtual screen) by protein-ligand docking was performed using Schrödinger Glide software.

### EN7 inhibition of TTLL4 in human AML cell lines

Powdered EN7 molecule was dissolved in cell-culture grade DMSO to a stock solution of 25 mM. Immediately prior to cellular treatment, the required volume of EN7 stock was diluted into 200 μL of fetal bovine serum (FBS) to facilitate solubility, mixed thoroughly and added to the respective media containing human AML cells. Control cells were treated with DMSO similarly diluted in FBS to a final concentration of 0.1%. CellTiter-Glo^®^ proliferation assays were performed using the manufacturer’s protocol (Promega) with the first day of treatment as day 0. Cells were collected for downstream experiments on days 3 and 6 of treatment, while remaining cells were passaged and resuspended in fresh media containing EN7 molecule prepared in FBS.

### Statistical analysis

GraphPad Prism 10 software was used for all statistical analyses. Specific statistical tests used for each analysis are indicated in the figure legends. Error bars indicate mean ± SEM. P values less than 0.05 were considered significant.

## Accession Numbers

RNA Sequencing data was deposited at the Gene Expression Omnibus under accession number GSE293535.

## Data Analysis Code

Code used in processing transcriptomic data can be found on GitHub:

https://github.com/Shechterlab/Schurer_Ilyas_etal_2025

## Acknowledgments

We would like to thank all past and current members of the Gritsman and Shechter laboratories for their helpful suggestions and discussions. We thank Simone Sidoli for assistance with mass spectrometry sample preparation. We thank the AECOM Flow Cytometry Core Facility (supported by NIH shared instrument grants 1S10OD026833-01 and 1S10OD032169-01) for assistance with flow cytometry and cell sorting. We thank Peter Schultes and Jason Justiniano of the AECOM Department of Cell Biology and also Simone Sidoli for technical assistance and facilities support. This work was supported by the Department of Defense Congressionally Directed Medical Research Program Peer Reviewed Cancer Research Program (CA200503) to D.S. and K.G., by the National Institutes of Health (R01GM135614) to D.S. and (R35GM155249) to S.K., by university seed funds to K.G, and by the Albert Einstein College of Medicine Venture Capital Pitch Prize to D.S., K.G., and S.K. D.S. was also supported by the Irma T. Hirschl Trust Career Scientist Award. Pilot funding to D.S. and K.G. was supported by the Montefiore Einstein Comprehensive Cancer Center under NCI Cancer Center Support Grant (P30CA013330). A.S. and M.I.M were supported by National Institutes of Health MSTP Training Grant (5T32GM007288-50). D.F.H. was supported by National Institutes of Health (NIH) grant GM037537. Experiment schematics were created using Biorender.com.

Data in this paper are from a thesis to be submitted in partial fulfillment of the requirements for the Degree of Doctor of Philosophy in the Biomedical Sciences at the Albert Einstein College of Medicine. The content is solely the responsibility of the authors and does not necessarily represent the official views of the National Institutes of Health.

## Author contributions

**A.S**. Conceptualization, resources, data curation, formal analysis, supervision, validation, investigation, visualization, methodology, writing–original draft, project administration, writing–review and editing. **H.I**. Conceptualization, resources, data curation, formal analysis, supervision, validation, investigation, visualization, methodology, project administration, writing–review and editing. **M.I.M**. Conceptualization, formal analysis, investigation, visualization, methodology, writing–review and editing. **S.H**. Investigation, visualization, methodology. **M.R.L**. Investigation, visualization, methodology. **I.R**. Investigation. **R.H**. Investigation. **L.D**. Investigation. **J.S**. Investigation. **D.F.H**. Investigation, methodology. **E.A**. Investigation, visualization. **V.M**. Investigation. **B.M.L**. Resources, validation. **S.G.S**. Investigation. **D.K.B**. Methodology. **Y.W**. Investigation. **L.M**. Resources, methodology. **R.B**. Resources. **S.K**. Funding, investigation, and analysis. **K.G**. Conceptualization, funding, analysis, visualization, writing-review and editing, and project supervision. **D.S**. Conceptualization, funding, analysis, visualization, writing-review and editing, writing–original draft, manuscript preparation, and project supervision.

