## Supplementary Figures S1-S7 for "TTLL4 glutamyltransferase is a therapeutic target for NPM1-mutated acute myeloid leukemia"

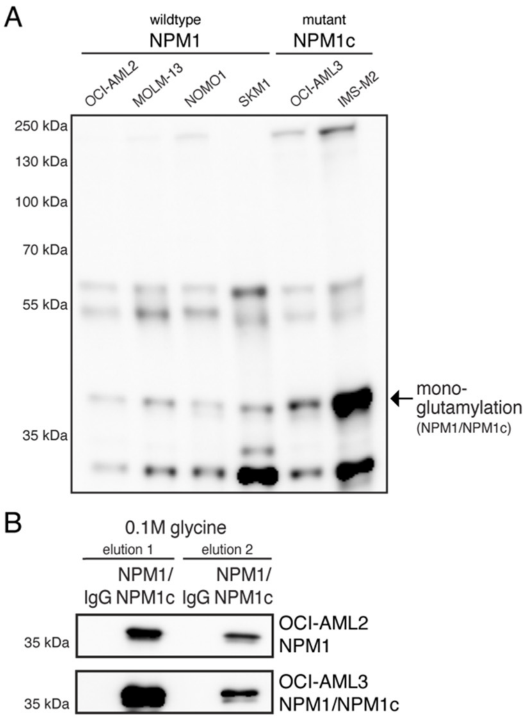


C

| sample | protein | E126 modification | total abundance of E126glu | E126glu with adjacent S125ph |
|---|---|---|---|---|
| recombinant protein | NPM1 | +1 glu | 22% | - |
| recombinant protein | NPM1c | +1 glu | 48% | - |
| OCI-AML2 | NPM1 | +1 glu | 10% | 100% |
| OCI-AML3 | NPM1 + NPM1c | +1 glu | 25% | 100% |
| IMS-M2 | NPM1 + NPM1c | +1 glu | 34% | 84% |

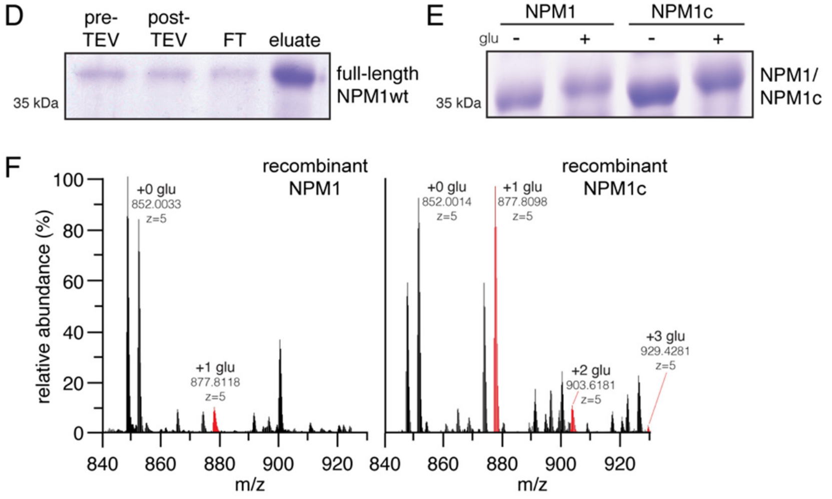


**Supplementary Figure S1. NPM1 and NPM1c are post-translationally glutamylated by TTLL4**
**(A)** Uncut immunoblot of mono-glutamylation in NPM1-wildtype and NPM1c-mutant AML cell lines. Arrow indicates ~37 kDa band corresponding to mono-glutamylated NPM1/NPM1c. **(B)** Immunoblots stained with total NPM1/NPM1c antibody depicting NPM1 and NPM1c proteins isolated from OCI-AML2 and OCI-AML3 cells using immunoprecipitation followed by glycine-mediated elution. These cellular proteins were also used for mass spectrometric analysis. **(C)** Table of quantification of percent abundance of E126 mono-glutamylated protein (E126glu) relative to unmodified protein and fraction of E126glu with phosphorylation of the adjacent serine 125 residue (S125ph) across the recombinant samples and three cell lines. **(D)** Coomassie blue stain depicting pre- and post-TEV cleaved variants of recombinant wildtype NPM1 protein. FT = flow-through. **(E)** Coomassie blue stain depicting the in vitro glutamylated and non-glutamylated variants of recombinant wildtype NPM1 and mutant NPM1c proteins used for mass spectrometric analysis. **(F)** Mass spectrometry (MS1) of glutamylation of recombinant NPM1 and NPM1c. Peaks are labeled with the C12 monoisotopic mass of each peptide.

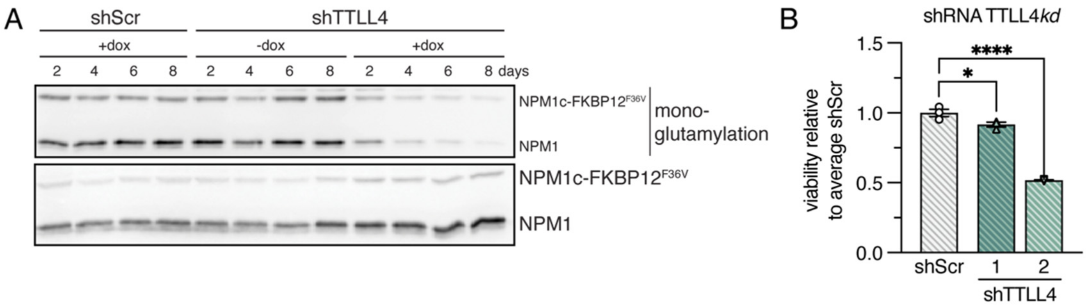


**Supplementary Figure S2. Loss of TTLL4 impairs proliferation and promotes myeloid differentiation of NPM1c-mutated AML cell lines**

Assays of NPM1c-degron OCI-AML3 cells with or without doxycycline-induced expression of shRNAs after lentiviral transduction of *TTLL4* (shTTLL4) or scrambled (shScr) shRNAs into NPM1c-FKBP12$^{F36V}$ (NPM1c-degron) OCI-AML3 cells. (A) Validation of shRNA-mediated *TTLL4* knockdown after doxycycline treatment by immunoblot analysis of mono-glutamylation of NPM1 (~37 kDa) and NPM1c-degron (~55 kDa). Analysis of NPM1 and NPM1c expression were used as loading controls. (B) Normalized cell viability at day 6 measured by CellTiter-Glo® proliferation assays. Error bars indicate mean ± SEM; *p ≤0.05, ****p ≤0.0001; ordinary one-way ANOVA.

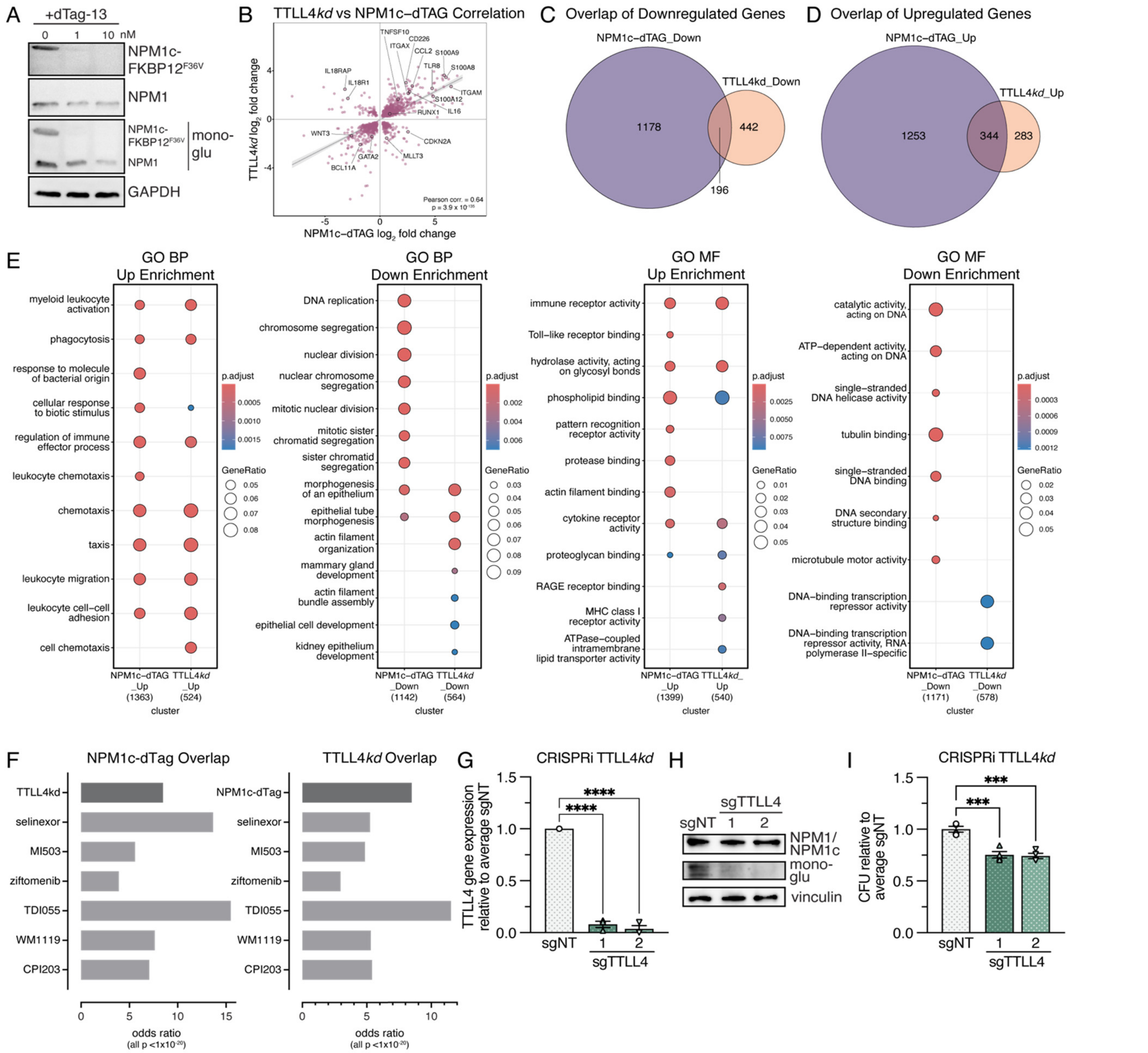


**Supplementary Figure S3. TTLL4 loss re-localizes NPM1/NPM1c to the nucleus and alters gene expression to promote myeloid differentiation**

(A) Immunoblot analysis assessing the expression of NPM1c-FKBP12F36V (~55 kDa), wildtype NPM1 (~37 kDa), and their mono-glutamylation following treatment with increasing doses of dTag-13. GAPDH was used as a loading control. (B) Scatterplot of genes significantly altered (padj < 0.01) in both NPM1c dTAG (x-axis) and TTLL4 knockdown (y-axis) showed a high and significant correlation (Pearson correlation = 0.64, p < 10-134). The same representative AML and differentiation markers are indicated as in Figures 3A and 3B. (C) Significantly altered (padj < 0.01) up-regulated genes in OCI-AML3 NPM1c-degron cell dTAG treatment mRNA seq were intersected with up-regulated genes in the TTLL4 knockdown and plotted in a Venn diagram. (D) Significantly altered (padj < 0.01) down-regulated genes in OCI-AML3 NPM1c-degron cell dTAG treatment mRNA seq were intersected with down-regulated genes in the TTLL4 knockdown and plotted in a Venn diagram. (E) Up- and down-regulated genes in both conditions were probed by Cluster-Profiler over-representation analysis for GO Biological Process (BP) and Molecular Function (MF) terms and plotted. (F) Significantly altered (padj < 0.01) genes in dTAG and TTLL4kd transcriptomes were compared with GeneOverlap to publicly available transcriptomes from OCI-AML3 cells treated with selinexor (72 hrs), MI503 and ziftomenib for menin inhibition, TDI055 for MLLT1 (ENL) inhibition, WM1119 for KAT6A acetyltransferase inhibition, and CPI203 as an iBET inhibitor. Odds ratios of overlap are shown, all Fisher Exact test p-values were < 10-20. (G) Validation of CRISPRi knockdown of TTLL4 in OCI-AML3 cells at day 10 by qPCR analysis of TTLL4 gene expression. (H) Validation of CRISPRi knockdown of TTLL4 in OCI-AML3 cells at day 10 by immunoblot analysis of mono-glutamylation of NPM1/NPM1c. Total NPM1/NPM1c expression and vinculin were used as loading controls. (I) Normalized number of colony-forming units (CFU) formed from 1 x104 cells by day 7 after CRISPRi knockdown of TTLL4. Error bars indicate mean ± SEM; ***p ≤0.001, ****p ≤0.0001; ordinary one-way ANOVA.

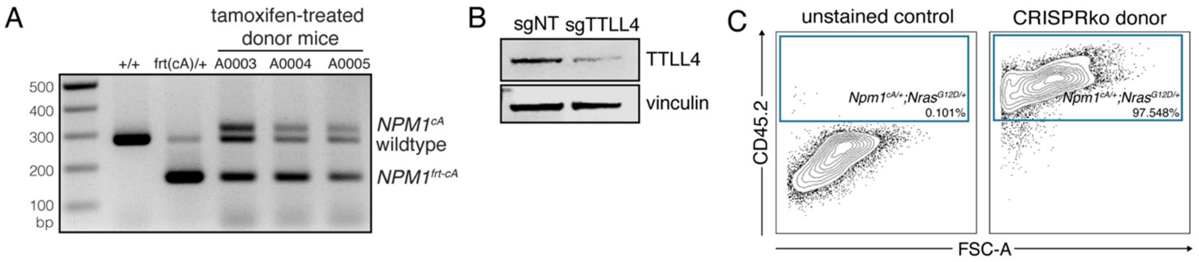


D

peripheral blood
5w post-transplantation

| group | mouse ID | WBC (K/uL) | Hb (g/dL) | PLT (K/uL) | TTLL4 indels (%) | editing analysis $R^2$ |
|---|---|---|---|---|---|---|
| sgNT (n=9) | A0207 | 50.98 | 5.8 | 463 | 0 | 1.00 |
| | A0208 | 56.56 | 9.5 | 390 | 0 | 1.00 |
| | A0209 | 18.70 | 8.8 | 348 | 0 | 1.00 |
| | A0210 | 48.18 | 9.9 | 458 | 0 | 1.00 |
| | A0212 | 39.16 | 14.1 | 363 | N/A | N/A |
| | A0213 | 54.78 | 12.6 | 418 | 0 | 1.00 |
| | A0214 | 36.62 | 11.2 | 497 | 0 | 1.00 |
| | A0215 | 42.38 | 12.8 | 265 | 0 | 1.00 |
| | A0216 | 29.98 | 11.1 | 303 | 0 | 1.00 |
| sgTTLL4 (n=9) | A0218 | 58.54 | 8.4 | 449 | 29 | 0.91 |
| | A0219 | 54.84 | 9.3 | 689 | 100 | 0.53 |
| | A0220 | 59.10 | 11.4 | 528 | 17 | 0.97 |
| | A0221 | 31.18 | 9.9 | 549 | 15 | 0.94 |
| | A0222 | 11.44 | 10.6 | 809 | 21 | 0.95 |
| | A0223 | 56.28 | 12.1 | 374 | 24 | 0.95 |
| | A0224 | 21.88 | 10.9 | 1467 | 25 | 0.95 |
| | A0225 | 12.86 | 11.2 | 519 | 100 | 0.50 |
| | A0226 | 36.20 | 12.1 | 530 | 29 | 0.89 |

**Supplementary Figure S4. TTLL4 is necessary for leukemia maintenance in mouse models of NPM1c AML**
(A) Validation of tamoxifen induction of heterozygous expression of the *Npm1$^{cA}$* allele in mouse peripheral blood cells. A0003, A0004, and A0005 were used as donors for CFU assays described in Fig. 4B. (B) Validation of CRISPR-Cas9 *Ttll4* deletion in sgTTLL4-RNP-electroporated *Npm1$^{cA/+}$* mouse bone marrow by immunoblot analysis of TTLL4 protein expression (130 kDa) with vinculin loading control. (C) Flow cytometry validation of high proportion of CD45.2$^{+}$ *Npm1$^{cA/+}$;Nras$^{G12D/+}$* cells in whole bone marrow used as donor cells for transplant experiments described in Fig. 4 and Fig. 6. (D) Analysis of peripheral blood of sgNT and sgTTLL4 *Npm1$^{cA/+}$;Nras$^{G12D/+}$* transplant recipient mice at 5 weeks post-transplantation, including white blood cell (WBC) counts, hemoglobin (Hb) levels, and platelet (PLT) counts. *Ttll4* CRISPR-Cas9 editing efficiency, represented by *Ttll4* indels percentage, was measured by Sanger sequencing and ICE analysis of DNA isolated from peripheral blood cells. The $R^2$ value indicates how well the indel distribution fits the sequence data of the sample.

A

bone marrow
survival endpoint

| group | mouse ID | survival (days) | liver weight (g) | spleen weight (g) | WBC (K/uL) | Hb (g/dL) | PLT (K/uL) | engraftment (% CD45.2+) | TTLL4 indels (%) | editing analysis $R^2$ |
|---|---|---|---|---|---|---|---|---|---|---|
| sgNT (n=9) | A0207 | 43 | 1.4659 | 0.4893 | 112.18 | 11.5 | 292 | 63.422 | 0 | 1.00 |
| | A0208 | 36 | 0.9842 | 0.2476 | 43.06 | 7.6 | 349 | 72.817 | 0 | 1.00 |
| | A0209 | 50 | 1.5654 | 0.3107 | 91.20 | 12.4 | 590 | 76.773 | 0 | 1.00 |
| | A0210 | 35 | 1.1343 | 0.2816 | 48.18 | 9.9 | 458 | 85.298 | 0 | 1.00 |
| | A0212 | 36 | 1.0524 | 0.2568 | 17.02 | 7.7 | 275 | 68.371 | 0 | 1.00 |
| | A0213 | 36 | 1.2258 | 0.3032 | 52.38 | 9.1 | 347 | 70.923 | 0 | 1.00 |
| | A0214 | 41 | 1.1580 | 0.2849 | 49.22 | 13.5 | 391 | 62.468 | 0 | 1.00 |
| | A0215 | 35 | 0.7735 | 0.2064 | 42.38 | 12.8 | 265 | 78.817 | 0 | 0.99 |
| | A0216 | 36 | 0.9473 | 0.2477 | 47.44 | 8.7 | 261 | 70.967 | 0 | 1.00 |
| sgTTLL4 (n=9) | A0218 | 52 | 1.5268 | 0.4547 | 169.12 | 8.5 | 337 | 59.098 | N/A | NA |
| | A0219 | 43 | 1.0061 | 0.3691 | 100.06 | 11.6 | 211 | 79.856 | 38 | 0.94 |
| | A0220 | 87 | 1.7107 | 0.6051 | 100.92 | 4..8 | 267 | 75.679 | 36 | 0.93 |
| | A0221 | 65 | 1.7043 | 0.4475 | 104.06 | 8.0 | 523 | 52.841 | 64 | 0.76 |
| | A0222 | 72 | 3.7805 | 0.9154 | 277.60 | 10.5 | 469 | 42.999 | 0 | 1 |
| | A0223 | 72 | 4.6446 | 0.6226 | 295.60 | 9.0 | 407 | 49.465 | 36 | 0.21 |
| | A0224 | 59 | 1.2442 | 0.3305 | 88.30 | 11.5 | 405 | 67.802 | 84 | 0.77 |
| | A0225 | 41 | 1.3010 | 0.2522 | 31.38 | 13.7 | 274 | 67.048 | 28 | 0.96 |
| | A0226 | 41 | 1.3193 | 0.3067 | 62.00 | 14.9 | 311 | 77.635 | N/A | N/A |

B

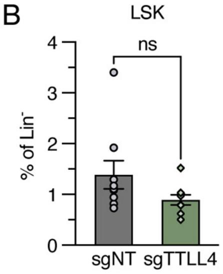


**Supplementary Figure S5. Loss of TTLL4 prolongs survival and promotes myeloid differentiation in NPM1c AML with a cooperating NRAS mutation.**
(A) AML disease burden analysis of sgNT and sgTTLL4 *Npm1*$^{cA/+}$;*Nras*$^{G12D/+}$ transplant recipient mice at survival endpoint, including liver and spleen weights and peripheral blood white blood cell (WBC) counts, hemoglobin (Hb) levels, and platelet (PLT) counts. Engraftment of *Npm1*$^{cA/+}$;*Nras*$^{G12D/+}$ donor cells was measured by flow cytometry quantification of the fraction of CD45.2$^+$ bone marrow cells. *Ttll4* CRISPR-Cas9 editing efficiency, represented by *Ttll4* indel percentage, was measured by Sanger sequencing and ICE analysis of DNA isolated from whole bone marrow. The $R^2$ value indicates how well the indel distribution fits the sequence data of the sample. (B) Quantification of flow cytometry analysis of the Lin$^-$ Sca1$^+$ c-Kit$^+$ (LSK) immature population in control (sgNT) or *Ttll4*-deleted (sgTTLL4) CD45.2$^+$ bone marrow collected at time of death. Representative flow plots are depicted in figure 4B. Error bars indicate mean ± SEM; unpaired t-test with Welch's correction.

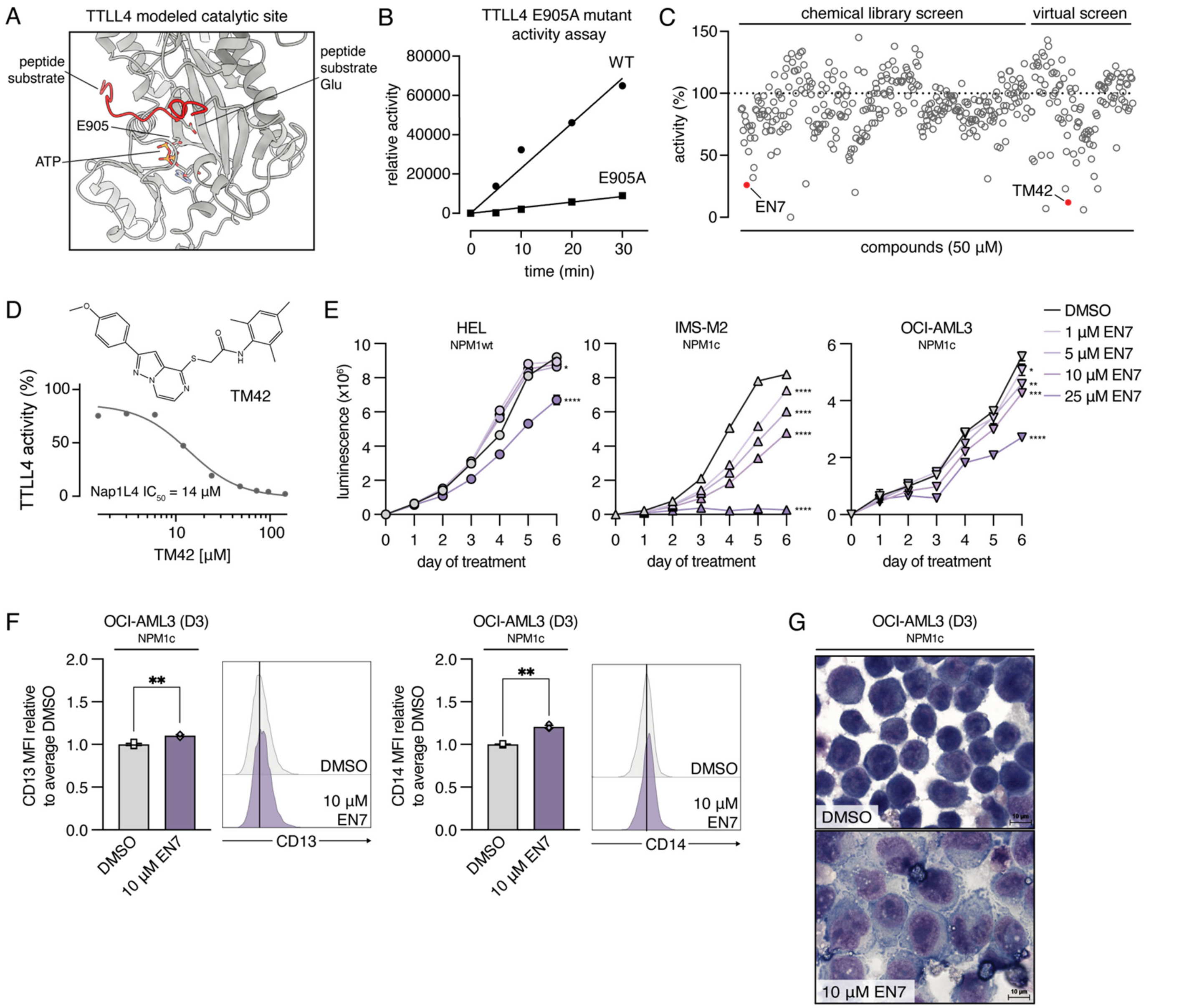


**Supplementary Figure S6. TTLL4 can be pharmacologically targeted in NPM1c AML**
(A) Modeled catalytic site of TTLL4 revealed that E905 was a putative critical residue for activity. (B) TTLL4 E905A mutant exhibited minimal glutamyltransferase activity compared with the control wildtype protein. (C) Combined screening data (50 µM molecule) from a 340-compound screen and from 114 candidate compounds from our computational virtual screen, incubated with TTLL4 and Nap1L4 substrate, are represented as percent of control activity. Compounds exhibiting loss of more than 50% of TTLL4 activity were further analyzed. EN7 and TM42 (red) were selected based on solubility and chemical properties for further study. (D) TM42 inhibited recombinant TTLL4 glutamyltransferase activity towards NAP1L4 with an *in vitro* $IC_{50}$ of 14 µM. (E) CellTiter-Glo® proliferation assays in NPM1-wildtype (HEL) and NPM1c-mutated (IMS-M2, OCI-AML3) human AML cell lines during treatment with increasing doses of EN7 or DMSO control. (F) Normalized median fluorescent intensity (MFI) of CD13 (left) and CD14 (right) expression in OCI-AML3 cells on day 3 of treatment with 10 µM EN7 or DMSO control. Representative histograms for one replicate are shown. (G) Wright-Giemsa staining of cytospins of OCI-AML3 cells on day 3 of treatment with 10 µM EN7 or DMSO control. Representative images from one experimental replicate are shown. Error bars indicate mean ± SEM; *p ≤0.05, **p ≤0.01, ***p ≤0.001, ****p ≤0.0001; ordinary two-way ANOVA (E), unpaired t-test with Welch's correction (F).

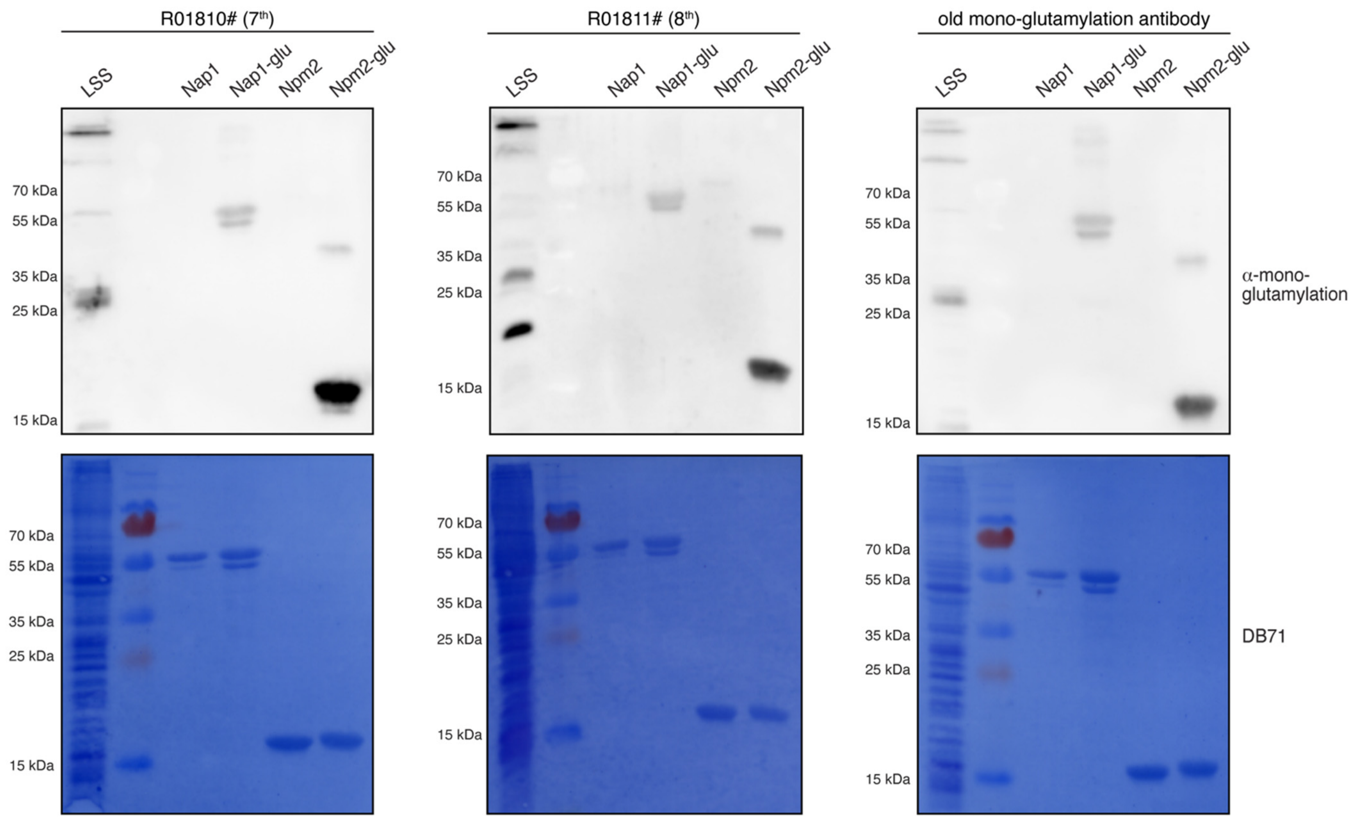


**Supplementary Figure S7. Validation of TTβIII/α-mono-glutamylation antibody**
Immunoblot analysis comparing the newly generated rabbit polyclonal mono-glutamylation antibodies (Rabbits #R10810 and R10811, following 7th and 8th immunizations, respectively) with a previously characterized antibody ("old"). Recombinant Nap1 and NPM2 proteins were untreated or enzymatically glutamylated *in vitro* with TTLL4, as indicated ("glu"). Direct Blue 71 (DB71) membrane staining was used as a loading control. The new antibodies specifically detect glutamylated Nap1 and NPM2, with minimal background reactivity, demonstrating improved specificity and sensitivity compared to the original antibody.
